# In vivo screening of tumor-hepatocyte interactions identifies Plexin B2 as a gatekeeper of liver metastasis

**DOI:** 10.1101/2023.05.15.540681

**Authors:** Costanza Borrelli, Morgan Roberts, Davide Eletto, Atefeh Lafzi, Jonas A. Kretz, Hassan Fazilaty, Marie-Didiée Hussherr, Elena Guido Vinzoni, Kristina Handler, Jan Michler, Srivathsan Adivarahan, Salvatore Piscuoglio, Xenia Ficht, Andreas E. Moor

## Abstract

It is estimated that only 0.02% of disseminated tumor cells are able to seed overt metastases^1^. While this indicates the presence of environmental constraints to metastatic seeding, the landscape of host factors controlling this process remains largely unknown. Combining transposon technology^2^ and fluorescent niche labeling^3^, we developed an *in vivo* CRISPR activation screen to systematically investigate the influence of hepatocytes on metastatic seeding in the liver. Our approach enabled the identification of Plexin B2 as a critical host-derived regulator of metastasis. Plexin B2 upregulation in hepatocytes dramatically enhances grafting in colorectal and pancreatic cancer syngeneic models, and promotes seeding and survival of patient-derived organoids. Notably, ablation of Plexin B2 in hepatocytes prevents mesenchymal-to-epithelial transition of extravasated tumor cells and thereby almost entirely suppresses liver metastasis. We dissect a mechanism by which Plexin B2 interacts with class 4 semaphorins on tumor cells, activating Rac1 signaling and actin cytoskeleton remodeling, thereby promoting the acquisition of epithelial traits. Our findings highlight the essential role of signals from the liver parenchyma for the survival of disseminated tumor cells, prior to the establishment of a growth promoting niche. They further suggest that acquisition of epithelial traits is required for the adaptation of extravasated cells to their new tissue environment. Targeting of Plexin B2 on hepatocytes shields the liver from colonizing cells and thus presents an innovative therapeutic strategy for preventing metastasis. Finally, our screening technology, which evaluates host-derived extrinsic signals rather than tumor-intrinsic factors for their ability to promote metastatic seeding, is broadly applicable and lays a framework for the screening of environmental constraints on metastasis in other organs and cancer types.

## Introduction

Historically viewed as a late-stage event in cancer progression, spreading of disseminated tumor cells (DTCs) has recently gained recognition as an early phenomenon in tumorigenesis^4–6^. Indeed, DTCs have been detected in patients with early breast cancer lesions^6^ and clinically undetectable colorectal cancer^7^. However, it is estimated that only 0.02% of DTCs are able to form overt metastases^1^, indicating major environmental barriers to metastatic seeding. Recent studies have elucidated mechanisms that promote DTC survival in circulation^8–10^, regulate DTC dormancy^11–13^, or promote recurrence^14^. Yet, due to the technical challenges of tracking single extravasated tumor cells, the environmental determinants of DTC adaptation and survival in a foreign organ environment remain largely unknown. Identification of these factors might reveal a therapeutic window to target metastasis at its most vulnerable point: prior to the establishment of a growth-promoting metastatic niche. To this end, we devised an experimental approach that enables in vivo screening of host organ-derived factors for their ability to regulate DTCs fate at the time of seeding. We applied it to uncover the hepatocyte-derived signals that regulate liver metastases. Our study identifies Plexin B2 as a crucial regulator of metastasis, revealing that its upregulation in hepatocytes dramatically enhances DTC seeding while its ablation suppresses liver metastasis by preventing mesenchymal-to-epithelial transition.

## Main

### Screening tumor-hepatocyte interactions in a mosaic liver

Hepatocytes constitute 60% of the liver by cell number and 80% by mass ^15^. We therefore hypothesized that early interactions with hepatocytes might influence the ability of extravasated DTCs to seed metastases. To test this, we developed an experimental strategy for pooled perturbation of hepatocyte-tumor interactions during seeding (**Fig. 1a**). First, hundreds of genes are stably overexpressed in hepatocytes using CRISPR-mediated transcriptional activation (CRISPR-a), resulting in a “mosaic liver” containing multiple perturbed seeding environments. Upon cancer inoculation, DTCs extravasating in the proximity of hepatocytes overexpressing a seeding-promoting factor will seed and grow, while DTC interacting with hepatocytes overexpressing a seeding-suppressing factor will fail to form a metastasis. A perturbation’s effect on seeding can therefore be inferred by its enrichment in hepatocytes neighboring successfully seeded metastases, indicating a promoting effect, or its enrichment in non-metastatic areas, indicating a seeding-suppressing effect (**Fig. 1a**). Thus, even if perturbed seeding events are not directly observed, their outcome can be retrospectively assessed by the presence, or absence, of a metastasis.

**Figure 1.**
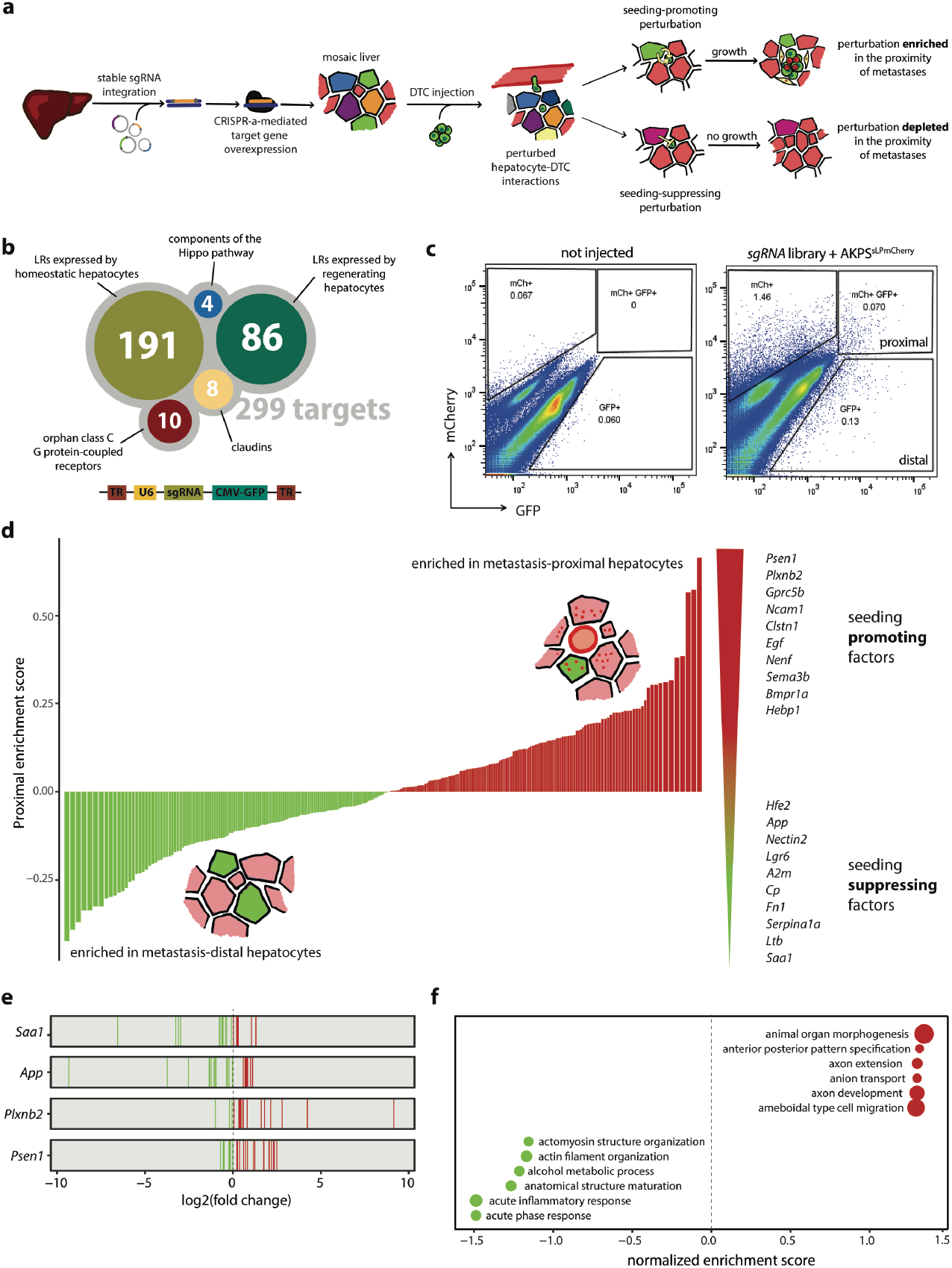
Screening tumor-hepatocyte interactions in a mosaic liver. **a**, Schematic representation of CRISPR-screening approach. Within the mosaic liver, extravasated tumor cells interact with hepatocytes harboring seeding-promoting or seeding-suppressing perturbations, resulting in successfull or failed metastatic outgrowth, respectively. **b**, Top, Genes targeted by the sgRNA library. Complete list in Table S1. Bottom, schematic of sgRNA vector design. **c**, Representative flow cytometry plots and gating strategy for the isolation of hepatocytes harboring a sgRNA (GFP^+^) and are either proximal to (mCherry^+^) or distal from (mCherry^-^) metastases. Hepatocytes gated as CD31^-^CD45^-^. **d**, Gene targets ranked by proximal enrichment score. Genes encoding for putative seeding-promoting factors are enriched GFP^+^mCherry^+^ hepatocytes in the proximity of metastases (red). Genes encoding for putative seeding-suppressing factors are enriched in GFP^+^mCherry^-^ hepatocytes in non-metastatic areas (green). Top-scoring putative seeding-suppressing and promoting factors listed on the right. **e**, logFC of individual sgRNAs (represented as vertical lines) for top-scoring suppressing and promoting factors across all mice and library batches. **f**, Gene set enrichment analysis of seeding promoting and suppressing factors. Dot size indicates the size of the gene set.

To achieve hepatocyte-specific expression of the CRISPR-a machinery, we crossed dCas9-SunTag-p65-HSF1^16^ with Albumin-Cre mice^17^. AlbCre;dCas9-SPH mice were then subjected to hydrodynamic tail vein injection (HTVI) of transposon plasmids harboring sgRNAs and Sleeping Beauty transposase (SB100X), which mediates stable integration and expression of exogenous DNA in mouse hepatocytes^2,18,19^. HTVI of sgRNAs targeting glutamine synthetase (GS, sgGlul), an enzyme normally only present in hepatocytes lining the central vein, resulted in ectopic and widespread GS immunoreactivity and increased Glul transcript expression (**Extended Data Fig. 1a, b**), indicating efficient on-target gene activation.

Molecular interactions between surface proteins expressed on tumor cells and hepatocytes are likely among the first events upon DTC extravasation into the space of Disse. We therefore designed a library of sgRNAs targeting ligands and receptors (LRs) genes expressed by murine hepatocytes^20^. Additionally, we included sgRNAs targeting LRs upregulated in the regenerating liver^20^; components of the Hippo signaling pathway, whose activation suppresses melanoma metastases via cell competition^21^; claudins, tight-junctions proteins recently implicated in CRC dissemination^22,23^; orphan class C G protein-coupled receptors (GPCRs) with unknown role in metastatic seeding^24^; and safe-targeting sgRNAs as negative controls^25^ (**Fig 1b**). The library (3 sgRNAs per gene, 997 sgRNAs in total, **Table S1)** was cloned into a transposon vector harboring CMV-GFP reporter expression, and co-injected with SB100X into AlbCre;dCas9-SPH mice at a concentration resulting in 0.5% GFP^+^ hepatocytes (**Extended Data Fig. 1c**). One week post-injection, we isolated CD31^-^CD45^-^GFP^+^ hepatocytes by fluorescent activated cell sorting (FACS) and performed targeted sgRNA amplification from genomic DNA followed by next generation sequencing. Correlation analysis of sgRNA abundances in the pre- and post-injection library reveals stable sgRNA distribution and high library retention in hepatocytes, indicating no loss of perturbation diversity (**Extended Data Fig. 1d**). Notably, the introduction of sgRNAs into non-proliferative hepatocytes, rather than into seeding tumor cells, ensures unaltered library distribution throughout the experiment, avoids bottleneck effects arising from poor grafting of tumor cells in vivo, and prevents library skewing towards perturbations that confer a proliferative advantage^26–28^. Moreover, the elevated number of hepatocytes in the adult murine liver (150 Mio^29^) enables screening of 997 sgRNAs library at high coverage (750x), while also ensuring a multiplicity of infection lower than 1 (MOI < 1).

Our approach depends on the assumption that a DTC interacting with a hepatocyte harboring a seeding-promoting perturbation would have increased chances of survival and therefore the corresponding sgRNA would be found enriched in the proximity of metastases. To record the proximity of perturbed hepatocytes to metastases, we employed a metastatic niche-labeling system based on a modified version of mCherry (sLP-mCherry) which is secreted by tumor cells and taken up by the neighboring cells^3^. We lentivirally introduced the sLP-mCherry reporter in Villin-CreER^T2^;APC^fl/fl^;Tp53^fl/fl^;Kras^G12D^;Smad4^KO^ (AKPS) organoids (AKPS^sLPmCherry^) and observed efficient peri-metastatic hepatocyte mCherry labeling in vivo upon intrasplenic injection (**Extended Data Fig. 1e, f, g**). Thus, hepatocytes with successful transposon insertion (GFP^+^) can be separated by fluorescence-activated cell sorting (FACS) as either proximal to metastasis (metastasis-proximal, mCherry^+^GFP^+^) or distant from metastases (metastasis-distal, mCherry^-^GFP^+^) (**Fig. 1c**).

We conducted three screening experiments with independently amplified sgRNA library batches (**Extended Data Fig. 2a**). In total, 7 AlbCre;dCas9-SPH mice and 5 nonCre littermate controls were injected with the sgRNA library and subsequently subjected to intrasplenic injection of AKPS^sLPmCherry^ organoids. Metastases were allowed to grow for 2 weeks, after which metastasis-proximal and distal hepatocytes were isolated by FACS. The amount of sorted cells across all experiments and mice resulted in cumulative coverage of 1000X for AlbCre;dCas9-SPH mice and 500X for the nonCre littermate controls (**Extended Data Fig. 2b, c**). We scored genes based on their proximal enrichment score, calculated based on the enrichment of their inferring sgRNAs in metastasis-proximal vs. distal hepatocytes (**Fig. 1d**). The highest scoring differentially enriched perturbations were consistent across individual mice and library batches (**Fig. 1e**), indicating high robustness of our screening strategy, and were not differentially enriched in nonCre littermates lacking dCas9-SPH expression (**Extended Data Fig. 2d**). Perturbations strongly enriched in metastasis-distal hepatocytes included several genes involved in acute phase reaction and immune response such as serum amyloid 1 (Saa1), amyloid precursor protein (App), and lymphotoxin-beta (Ltb) (**Fig. 1f, g**). The enrichment of sgRNAs targeting these genes in non-metastatic areas suggests that their upregulation prevents seeding of DTCs, possibly by acting as myeloid chemoattractants^30,31^ and inducing local immune activation. Indeed, Saa1 was suggested to attract macrophages to the tumor invasive front^30^, while amyloid protein (App) deposition recruits neutrophils in pancreatic, skin and lung cancer^31^. Conversely, overexpression of genes targeted by sgRNAs found enriched in the proximity of metastases induce, has favorable effect on DTCs seeding. These included Egf, a known driver of metastatic CRC^32,33^, as well as other regulators of morphogenesis (Gpc3, Psen1) (**Fig 1d-g**). Interestingly, our screen further implicated several LRs involved in axon guidance such as Plxnb2, Nrp2, Ncam1 and Nenf as seeding-promoting factors. This is in line with reports of DTCs hijacking axonal morphogenesis pathways to interact with endothelial cells ^34,35^, however their role in tumor-hepatocyte interactions has not been explored.

### Plexin B2 has a direct growth-promoting effect on tumor cells

We cross-validated the results of our screen by analyzing human transcriptional and mutational data. We first evaluated whether any of the putative seeding-regulating factors are predicted to interact with tumor cells. To this end, we generated spatial transcriptomics data (Visium, 10x Genomics) of hepatic CRC metastases and performed cell type deconvolution of spots using published single-cell RNA sequencing datasets ^36,37^ (**Extended Data Fig. 3a**). We then subdivided hepatocyte-containing spots in metastasis-proximal and metastasis-distal, and tumor cell-containing spots in metastasis edge and core, and used NicheNet^38^ to predict LR interactions between metastasis-proximal hepatocytes and tumor cells at the metastatic leading edge (**Fig. 2a**). Among the 109 predicted interactions, 22 involved top-scoring hits of our screen, including the chemoattractants App, Saa1 and Ltb and the axon guidance molecules Psen1, Plxnb2, Nenf and Nectin2.

**Figure 2.**
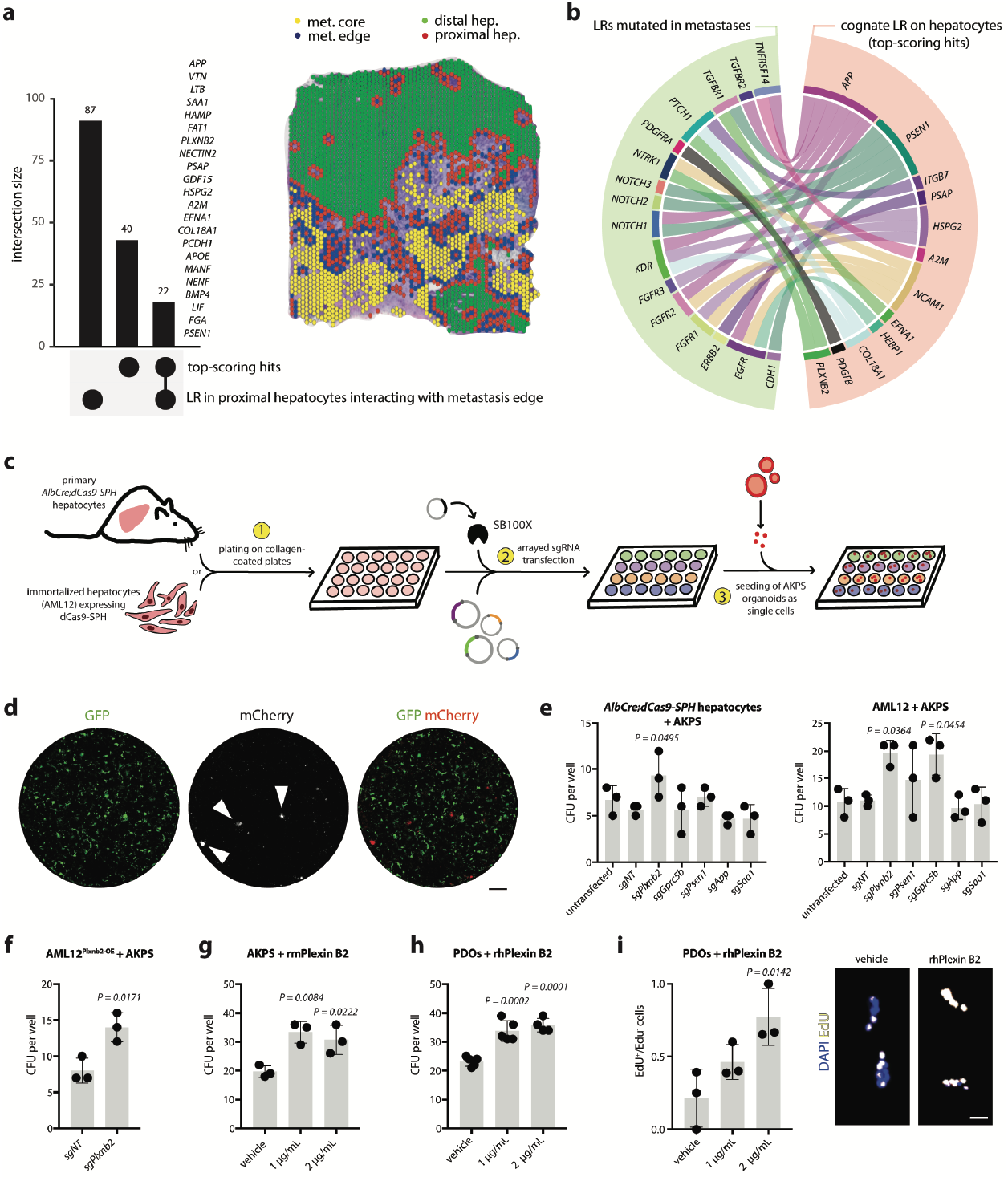
Plexin B2 has a direct growth-promoting effect on tumor cells. **a**, LR interaction analysis at metastatic leading edge. Upset plot showing the intersection of screen hits with LR genes involved in interactions between metastasis proximal hepatocytes (red spots) and tumor cells at metastatic edge (blue spots). Intersecting genes are listed. **b**, Circus plot of interactions between frequently mutated LRs in hepatic metastases and top-scoring hits expressed by hepatocytes. **c**, Experimental workflow for in vitro arrayed screen. 1) primary *AlbCre;dCas9-SPH* hepatocytes or *dCas9-SPH*-expressing AML12 cells are plated on collagen-coated plates and 2) transfected with arrayed sgRNA and SB100X, prior to 3) seeding of dissociated AKPS^sLP-mCherry^ organoids. After 5 days, CFUs are scored by fluorescent microscopy. **d**, Representative fluorescence micrographs showing transfected hepatocytes (GFP^+^) and AKPS colonies (mCherry^+^, indicated with arrowheads) after 5 days of co-culture. Scale bar, 100 μm. **e**, CFU per well of AKPS^sLP-mCherry^ organoids after co-culture with primary hepatocytes (left) or ALM12 cells (right) transfected with indicated sgRNAs. **f**, CFU per well of AKPS^sLP-mCherry^ organoids after co-culture with AML12 cells stably and uniformly overexpressing *Plxnb2* (AML12^Plxnb2-OE^). Two-tailed unpaired t-test. **g**, CFU per well of AKPS^sLP-mCherry^ organoids after co-culture with AML12 cells with increasing concentrations of rmPlexin B2. **h**, CFU per well of patient-derived CRC organoids (PDOs) cultured with rhPlexin B2. **i**, EdU incorporation PDOs cultured with rhPlxnb2 (left) and representative fluorescent micrograph of Edu+ PDO colonies (right). Scale bar, 20 μm. **e-i**, Barplots indicate mean ± SD. Dots represent individual wells. **c, e, g** Ordinary one-way ANOVA, comparing each treatment group to *sgNT* or vehicle control. Tukey’s correction for multiple testing.

We next wondered whether any seeding-regulating factors could interact with genes mutated in hepatic metastases, possibly indicating a selection of these interactions. To test this, we filtered a published dataset of genomic alterations more frequently found in hepatic metastases compared to matched primary tumors^39^ for genes encoding for ligands and receptors (**Extended Data Fig. 3b**). We then used LR interaction databases^38,40,41^ and GTEx data to identify their putative interaction partners expressed by hepatocytes. We found that 16 LRs frequently mutated in hepatic metastases were predicted to interact with top-scoring hits of our screen such as Plxnb2, Psen1, App, and Ncam1 (**Fig. 2b, Extended Data Fig. 3c**). Together with our screening results, these analyses implicate hepatocyte-derived chemoattractants and axon guidance cues as regulators of metastatic seeding in the liver (**Extended Data Fig. 3d**).

To test the direct influence of these factors on cancer growth, we devised a small plate-based interaction screen based on co-culture of hepatocytes and cancer cells. sgRNAs targeting top-scoring hits of our screen were transfected in an arrayed fashion in primary hepatocytes isolated from AlbCre;dCas9-SPH mice, or in immortalized murine hepatocytes (AML12) expressing dCas9-SPH. AKPS^sLP-mCherry^ organoids dissociated into single cells were then sparsely seeded on the hepatocyte monolayer and allowed to grow for 5 days before counting of colonies (CFU, colony forming units) (**Fig. 2c, d**). Overexpression of App, Saa1 did not result in significantly altered CFU with respect to non-targeting sgRNA (sgNT) or untransfected controls, suggesting that their effect on seeding might be mediated local recruitment of third cell type (**Fig. 2e**). In line with our in vivo results, we observed increased CFU upon Plnxb2, Psen1 and Gprc5b overexpression, indicating their direct involvement in the interaction between hepatocytes and tumor cells (**Fig. 2e**). In particular, overexpression of Plxnb2 had a very potent promoting effect on seeding. Single-cell seeding of AKPS organoids on AML12 cultures uniformly overexpressing Plxnb2 led to a 2-fold increase in CFU numbers (**Fig. 2f**). Moreover, addition of recombinant murine Plexin B2 ectodomain (rmPlexin B2) on AKPS single cells significantly increased CFU both in the presence and absence of hepatocytes, suggesting that Plexin B2 directly binds tumor cells (**Fig. 3g, Extended Data Fig. 3e**). Notably, these results could be recapitulated by adding recombinant human Plexin B2 (rhPlexin B2) on patient-derived CRC organoid (PDO) line seeded as single cells in the presence or absence of human immortalized hepatocytes (PTA-5565) (**Fig. 3h, fig. Extended Data Fig. 3f, g**). PDO colonies treated with rhPlexin B2 further exhibited increased numbers of EdU^+^ cells, indicating enhanced proliferation (**Fig. 3i**). These results prompted us to investigate the role of Plexin B2 in mediating interactions between hepatocytes and seeding DTCs in vivo.

**Figure 3.**
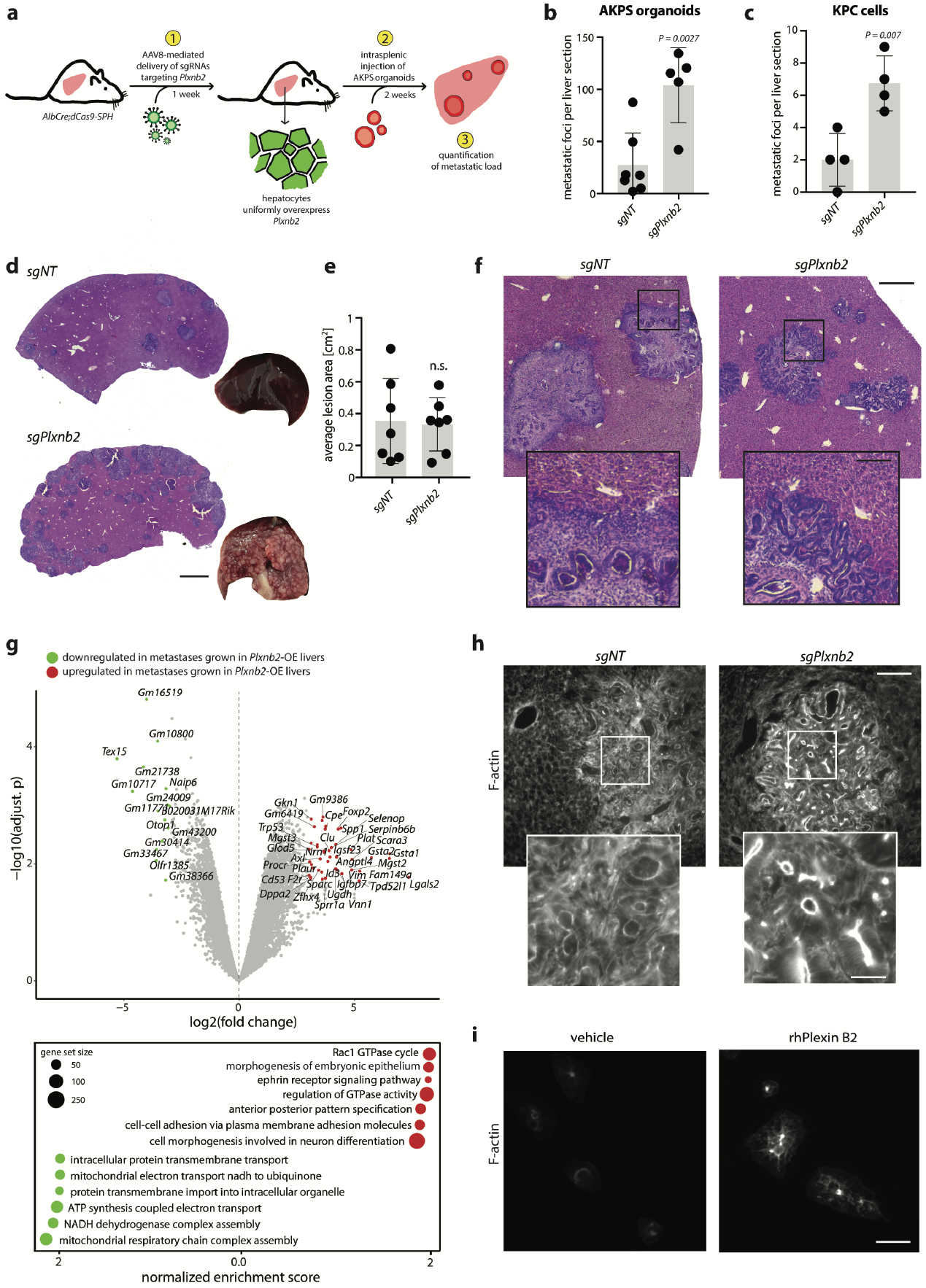
Overexpression of Plexin B2 promotes seeding of metastases in vivo. **a**, Schematics of experimental workflow. 1) *AAV8-U6-sgPlxnb2-EF1A-EGFP*-mediated overexpression of *Plxnb2* in hepatocytes is followed by 2) intrasplenic injection of AKPS organoids and 3) quantification of metastatic load. **b-c**, Quantification of metastatic foci per liver section 2 weeks after injection of AKPS organoids (b) or KPC cells (c). **d**, Representative H&E staining and pictures of livers harboring *Plxnb2*-overexpressing hepatocytes (*sgPlnxb2*) or control livers (*sgNT*) colonized by AKPS metastases. Scale bar, 500 μm. **e**, Average area in cm^2^ of AKPS metastases in individual sections of *sgNT* or *sgPlxnb2* livers. **f**, Representative H&E staining of AKPS metastases in sgNT or sgPlxnb2 livers. Scale bar, 200 μm. Insets show metastasis leading edge. Scale bar, 50 μm. **g**, Differentially expressed genes (top) and enriched GO-terms (bottom) in AKPS metastases grown in *Plxnb2*-overexpressing vs control livers. **h**, Representative fluorescent micrographs of F-actin staining in metastases grown in Plxnb2-overexpressing compared to control livers. Scale bar, 100 μm. Insets show F-actin accumulation at the apical side of epithelial duct-like structures. Scale bar, 20 μm. **I**, Representative fluorescent micrographs of F-actin staining of CRC PDOs in the presence or absence of rhPlexin B2. **b**,**c**,**e**, Barplots indicate mean ± SD. Dots represent individual mice, two-tailed unpaired t-test.

### Hepatocyte-derived Plexin B2 increases the rate of seeding events

Plexin B2 is widely expressed in the nervous system, as well as in epithelial cells of most mouse tissue, where it localizes to the basolateral membrane^42^. Its functions are mostly characterized in neural development^43^, however the phylogenetic emergence of the plexin family predates the appearance of the nervous system^44^ and recent studies have unraveled roles of Plexin B2 in several tissue contexts^42,45–48^. In order to investigate the seeding-promoting effect of hepatocyte-derived Plxnb2 overexpression in vivo, we used adeno-associated virus 8 (AAV8) to broadly deliver sgRNAs targeting Plxnb2 (AAV8-U6-sgPlxnb2-EF1A-EGFP) to hepatocytes in AlbCre;dCas9-SPH mice (**Extended Data Fig. 4a**). This induced a 2-fold upregulation of Plxnb2 expression in hepatocytes, as assessed by single molecule fluorescent in situ hybridization (smFISH) and Plexin B2 immunofluorescence (**Extended Data Fig. 4b, c, d**). In line with our screen, intrasplenic injection of AKPS organoids in livers harboring Plxnb2-overexpressing hepatocytes resulted in a 3-fold increase in metastatic foci compared to injections in control livers (**Fig. 3a, b**). Remarkably, Plxnb2 overexpression also increased grafting of syngeneic pancreatic ductal adenocarcinoma cells (Ptf1a-Cre;Kras^G12D/+^;Trp53^flox/+^, KPC), suggesting that its promoting effect applies to other cancers of epithelial origin that frequently metastasize to the liver (**Fig. 3c**). Interestingly, the average area of AKPS metastases in Plxnb2-overexpressing livers was unaltered with respect to those in control livers, indicating that Plexin B2 increases the rate of successful seeding, rather than conferring a proliferative advantage to growing metastases (**Fig. 3d, e**).

Histological analysis revealed that lesions in Plxnb2-overexpressing livers lacked the fibrous rim that usually surrounds AKPS metastases and prevents direct contact between tumor cells and hepatocytes (**Fig. 3f**). This morphology is reminiscent of the replacement-type histopathological growth pattern (HGP) of human CRC hepatic metastases, which lack extensive stromal remodeling and is associated with shorter overall survival and resistance to antiangiogenic treatment^49,50^. Metastases in Plxnb2-overexpressing livers indeed harbored decreased numbers of **α**-smooth muscle actin^+^ (α-SMA) and periostin^+^ (POSTN) cancer-associated fibroblasts (CAFs), and exhibited an extensive CD146^+^ vascular network (**Extended Data Fig. 4e**). Of note, α-SMA and POSTN immunoreactivity in Plxnb2-overexpressing livers was unaltered, suggesting that changes in the stromal and endothelial compartments were not induced prior to AKPS intrasplenic injection (**Extended Data Fig. 4g**). Moreover, ex vivo treatment of AKPS organoids with rmPlexin B2 was sufficient to increase their metastatic seeding in wild-type (WT) livers (**Extended Data Fig. 4h**). This, together with our in vitro results, indicates that Plexin B2 has a direct effect on the grafting of DTCs.

To assess the effect of overexpressing Plexin B2 on the tumor transcriptome, we profiled microdissected metastases grown in Plxnb2-overexpressing and control livers, and found upregulation of gene sets related to epithelial morphogenesis, Rac1 GTPase signaling, neuronal development, pattern specification and adhesion (**Fig. 3g**). Consistent with Rac1 activation and cytoskeletal remodeling, phalloidin staining indicated striking apical F-actin accumulation in duct-like structures of metastases grown in Plxnb2-overexpressing liver (**Fig. 3h**). Moreover, treatment of CRC PDOs with rhPlexin B2 in vitro induced F-actin focal aggregation (**Fig. 3i**). Together, these observations indicate that Plexin B2 on hepatocytes not only provides survival signals to extravasated AKPS cells - increasing the rate of successful seeding events - but also profoundly alters the state of growing lesions and their interactions with the recruited metastatic microenvironment.

### Plexin B2 induces mesenchymal-to-epithelial transition

Our results implicate hepatocyte-derived Plexin B2 in direct interaction with DTCs. Recently, therapies targeted at the tumor microenvironment, rather than tumor cells, have emerged as promising avenues to limit cancer progression^51^. We therefore wondered whether targeting Plexin B2 on hepatocytes could represent a therapeutic strategy to shield the liver from metastatic colonization. To test this, we used AAV8 injection to deliver sgRNAs targeting Plxnb2 in AlbCre;Cas9 mice, which resulted in hepatocyte-specific ablation of Plexin B2 (**Extended Data Fig. 5a)**. Remarkably, loss of Plexin B2 almost completely prevented metastatic outgrowth of AKPS organoids (**Fig. 4a, b**). The few remaining lesions in Plxnb2-KO livers were smaller in size, exhibited cellular disarray and mostly lacked duct-like structures, suggesting that absence of Plexin B2 impairs proliferation, epithelial morphogenesis and establishment of polarity (**Fig. 4c**).

**Figure 4.**
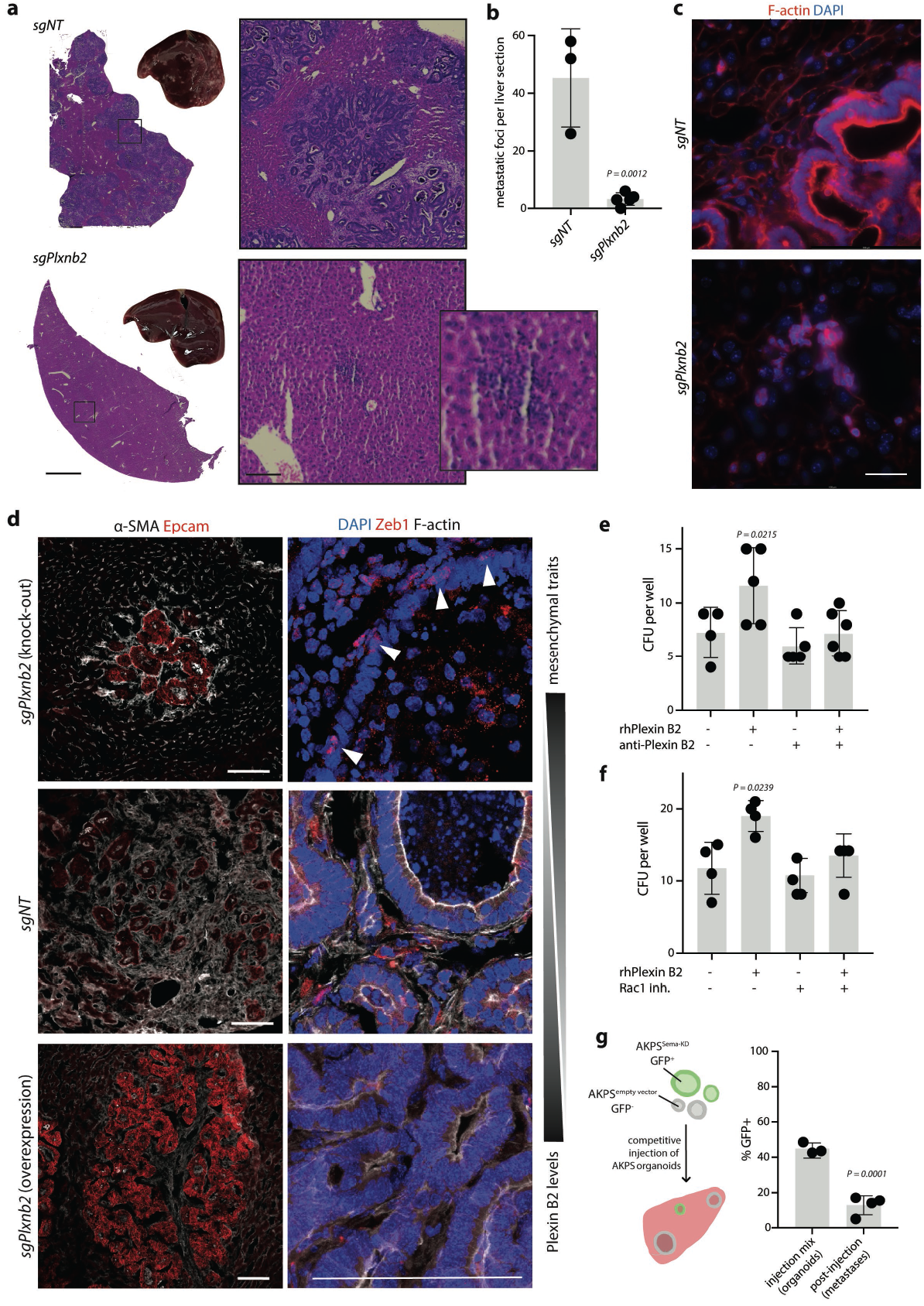
Plexin B2 is required for liver colonization. **a**, Representative H&E staining and pictures of livers harboring *Plxnb2*-KO hepatocytes (*sgPlnxb2*) or control livers (*sgNT*) colonized by AKPS metastases. Scale bar, 500 μm. Inset show detail of metastasis. Scale bar, 200 μm. **b**, Metastatic foci per liver section in *AlbCre*;*Cas9* mice harboring *Plxnb2*-KO (*sgPlxnb2*) or control hepatocytes (sgNT). **c**, Representative fluorescent micrographs of F-actin in AKPS metastases grown in *Plxnb2*-KO or control livers. Scale bar, 100 μm. **d**, Representative fluorescent micrographs of AKPS metastases grown Plxnb2 KO (*AAV8-sgPlxnb2* in *AlbCre;Cas9* mice), control (*AAV8-sgNT* in *AlbCre;Cas9* mice), or in Plxnb2 overexpressing livers (*AAV8-sgPlxnb2* in *AlbCre;dCas9-SPH* mice). Top, “-SMA (white) and Epcam (red) immunoreactivity. Bottom, nuclear DAPI (blue) and Zeb1 (red) immunoreactivity. Arrowheads show epithelial cells with nuclear Zeb1 localization. Scale bar, 100 μm. **e**, CFU per well of patient-derived CRC organoids (PDOs) with 2 !g/mL rhPlexin B2 and 1 ng/!L anti-Plexin B2 antibody targeting the semaphorin domain, or IgG1 isotype control. **f**, CFU per well of patient-derived CRC organoids (PDOs) cultured with 2 !g/mL rhPlexin B2 and 50 !M Rac1 inhibitor NSC23766. **g**, left, Schematic representation of competitive seeding assay. AKPS organoids harboring short hairpin (sh)RNAs against *Sema4a, Sema4c, Sema4d* and *Sema4g* (Sema-KD, marked by GFP) are mixed in a 1:1 ratio with control organoids (empty vector) and co-injected intrasplenically. After 2 weeks, the amount of GFP^+^ and GFP^-^ metastases is quantified. Right, Percentage of GFP^+^ metastases (post-injection) compared to percentage of GFP^+^ organoids (pre-injection). **b**,**e**,**f**,**g**, Barplots indicate mean ± SD. **b**,**g**, Dots represent individual mice, two-tailed unpaired t-test. **e, f**, Ordinary one-way ANOVA, comparing each treatment group to untreated control. Tukey’s correction for multiple testing.

The striking effect of Plexin B2 up- or downregulation on the morphology of AKPS metastases led us to hypothesize that it might promote mesenchymal-to-epithelial transition (MET) in extravasated DTCs. Reversion of EMT is thought to be essential for successful metastatic outgrowth of breast and squamous cell carcinomas^52,53^, however its involvement in the establishment of CRC metastases is largely unknown. While AKPS organoids in culture are in a hybrid EMT state, characterized by a mix of more epithelial (E-cadherin^high^Zeb1^low^) and more mesenchymal cells (E-cadherin^low^Zeb1^high^) (**Extended Data Fig. 5b**), liver metastases in WT and Plxnb2-overexpressing livers completely lack Zeb1 expression, indicating MET (**Fig. 4d**). We further observed increased Epcam immunoreactivity by metastases grown in Plxnb2-overexpressing livers, an additional similarity with human metastases of the replacement HGP, which exhibit increased EPCAM transcript levels compared to the desmoplastic HGP^50^. The upregulation of the transcription factors FoxP2 and Id3 upon Plxnb2 overexpression (**Fig. 3g**) is further consistent with repression of mesenchymal programs^54,55^. Strikingly, metastases in Plxnb2-KO livers retained epithelial Zeb1 expression (**Fig 4e**), indicating that absence of Plexin B2 on hepatocytes prevents suppression of mesenchymal programs. The minimal grafting of AKPS organoids in Plxnb2-KO livers suggests that Plexin B2-induced MET is required for DTC seeding and adaptation to the liver environment. Indeed, organoid lines derived from AKPS metastases grown in WT liver with physiological Plexin B2 levels exhibit morphological and transcriptomic evidence of epithelial differentiation, and have impaired ability to establish colonies when seeded in vitro as single cells, indicating increased susceptibility to anoikis^56^ (**Extended Data Fig. 5c-e**).

Canonical Plexin B2 ligands include class 4 semaphorins, which have been previously implicated in cancer cell growth, migration, invasive reprogramming, and metastasis formation^57–61^. In a large cohort of CRC patients, increased expression of SEMA4A, SEMA4C and SEMA4D expression was indeed associated with reduced recurrence-free survival (**Extended Data Fig. 6b**). Class 4 semaphorins have been reported to act both as ligands and as receptors, activating reverse signaling via Rac1, and suppressing mesenchymal programs^55,57,62^, suggesting that these ligands might mediate the seeding-promoting effects of Plexin B2 on DTCs. In line with this hypothesis, both antibody-mediated blockade of the semaphorin-binding domain of Plexin B2, and Rac1 inhibition prevented the seeding-promoting effect of rhPlexin B2 on PDOs in vitro (**Fig. 4e, f**). However, treatment with Pepinemab, a semaphorin 4D blocking antibody currently trialed for immuno-oncological and neurological diseases^63^, did not impair seeding of PDOs, hinting at redundant roles of the semaphorin paralogues in mediating Plexin B2 signaling in tumor cells (**Extended Data Fig. 6b**). Indeed, we detected expression multiple semaphorins reported to interact with Plexin B2, namely Sema4a, Sema4c, Sema4d, Sema4g, as well as Plxnb2 mRNA in AKPS metastases by multiplexed in situ hybridization (Molecular Cartography, **Extended Data Fig. 6c**). We therefore performed simultaneous partial knock-down of Sema4a, Sema4c, Sema4d, and Sema4g in AKPS organoids (Sema-KD), and observed downregulation of gene sets involved in epithelial morphogenesis, cell adhesion and Rac1 GTPase activity (**Extended Data Fig. 6d-f**). Notably, when co-injected with control organoids in competitive seeding assays, Sema-KD AKPS organoids exhibited reduced grafting ability, indicating that class 4 semaphorins are required for efficient liver colonization in vivo (**Fig. 4g, Extended Data Fig. 6g**).

Cumulatively, our results show that Plexin B2 is a host-organ derived factor that regulates the fate of seeding DTCs by inducing mesenchymal-to-epithelial transition. We show that Plexin B2 on hepatocytes directly interacts with semaphorins on tumor cells, profoundly altering their state and their interactions with the stromal and endothelial compartment. Our results further indicate that the semaphorin-plexin signaling, for example by triggering Plexin B2 internalization, might serve as a therapeutic target to prevent metastatic colonization of the liver.

## Discussion

The importance of microenvironmental conditions of the host organ for metastatic seeding has long been recognized. The seminal “seed and soil” hypothesis postulates that metastatic cells will only seed and colonize favorable environments^64^. However, we still lack a comprehensive overview of the signals from the metastasis-accepting organs that promote or suppress establishment of secondary tumors. Here, we devised an experimental approach for massively parallel in vivo screening of hepatocyte-derived factors that regulate metastatic seeding in the liver. Indeed, while they constitute the majority of the liver parenchyma, the role of this cell type in regulating metastases is largely unknown.

The data presented herein identify hepatocyte-derived Plexin B2 as required for liver colonization, and its upregulation sufficient to dramatically increase metastatic burden in both colorectal and pancreatic carcinoma models. Our results further suggest that Plexin B2 on hepatocytes is an essential inducer of MET in seeding metastases. The reversion of EMT is required for tumor cells with epithelial origin to survive, proliferate and form secondary tumors^52,53,65–68^, but the mechanisms regulating this switch are not well understood. Plexin-dependent semaphorin 4C reverse signaling has previously been linked to suppression of mesenchymal transcription factors in breast cancer^55^, while semaphorin 4A and 4D have been shown to control cancer cell migration through Rac1 activity^57,62^. Our results corroborate these findings and implicate reverse signaling from semaphorins as mediating the seeding-promoting effects of Plexin B2 in tumor cells. Yet, a fine mechanistic dissection of the molecular events downstream of the interaction between hepatocyte-derived Plexin B2 and tumor-derived semaphorins remains to be achieved.

Retrospective analysis may identify factors that allow metastases to thrive long-term^39,69–72^, but fails to capture early events of metastatic seeding, and distinguish between cellular interactions that are cause or consequence of metastatic outgrowth^3,73–75^. In contrast, our screening method allows for functional testing of cell-cell interactions in vivo, enabling the identification of those that regulate the fate of DTCs at the time of seeding. Importantly, our experimental approach can be broadly applied. We envision screening of other liver-resident cell types such as hepatic stellate cells and sinusoidal endothelial cells, which have been implicated in regulating liver colonization^76–78^. The system could also be adapted to perturb seeding events in other organs such as the lungs, lymph nodes, brain or bone marrow. We here focus on CRC, however, other cancer types that frequently disseminate to the liver such as melanoma, pancreatic, gastric or breast cancer^79^ might require different hepatocyte-derived seeding factors, and merit independent screening. Finally, additional libraries could be designed to probe environmental factors with a putative involvement in the early metastatic niche, such as extracellular matrix deposition^80,81^, degradation^82,83^ or metabolic pathways ^84–86^. In sum, the results presented herein identify Plexin B2 as a gatekeeper of liver metastasis, and lay a methodological framework to deepen our understanding of metastatic seeding.

## Supporting information

Supplemental Gating Strategy

## End notes

## Acknowledgements

We thank all members of the Moor lab, Gustavo Aguilar and Tomas Valenta for insightful discussions. We are also thankful to the Single Cell Facility of D-BSSE and to members of the laboratories of Konrad Basler, Maries van den Broek, Gerald Schwank and Randall Platt for technical support. We are grateful to Owen Sansom (Beatson Institute for Cancer Research, Glasgow) for donating the AKP organoids, to Mohamed Bentires-Alj (University of Basel) for providing the human immortalized hepatocytes, and to Ilaria Guccini (ETH Zurich) for donating the KPC cells and to Thorsten Stühmer for donating the shRNA plasmid backbone (University Hospital of Würzburg). This work was funded by the Swiss National Science Foundation (PCEFP3_181249 to A.E.M.) and the Swiss Cancer League (KFS-5444-08-2021-R to C.B. and A.E.M.). S.P. si funded by the Swiss Cancer Research foundation (KFS-4988-02-2020-R) and by The Prof. Dr. Max Cloëtta foundation. This paper was typeset with the bioRxiv word template by @Chrelli: www.github.com/chrelli/bioRxiv-word-template.

## Author contributions

C.B. and A.M. conceived the study and designed experiments. C.B., M.R., D.E., M.H.D., J.A.K, J.M. and K.H. performed the experiments. C.B., E.G.V, S.A., A.L. and K.H. performed the data analysis. S.P. obtained the PDOs. C.B. wrote the manuscript. X.F. contributed to editing and writing the manuscript. A.M. supervised the study.

## Materials and Methods

### Mice

All experiments were performed on 6-16 week-old male and female mice. Albumin-Cre mice (AlbCre, stock no. 003574), LSL-dCas9-SPH (dCas9-SPH, stock no. 031645), LSL-Cas9 (Cas9, stock no. 024858) and mT/mG mice (stock no. 007576) were obtained from a local live mouse repository. Chow and water were available ad libitum, unless specified. All mice were in the B6J background and maintained on a 12h light / 12h darkness schedule. Mice were housed and bred under specific pathogen-free conditions in accredited animal facilities. At the experimental endpoint, mice were sacrificed by raising CO_2_ concentrations. All experimental procedures were performed in accordance with Swiss Federal regulations and approved by the Cantonal Veterinary Office. Genotyping was performed by Transnetyx genotyping services.

### Animal experiments

Hydrodynamic tail vein injection: Hydrodynamic injection is an efficient procedure to deliver nucleic acids to the liver which involves the rapid injection (6-8 sec) of a large volume (8-12% body weight) of saline (0.9% sodium chloride) into the tail^87^. Mice were placed in a restraining device, and the tail was warmed with a red light lamp to induce lateral tail vein dilatation. The injection site was cleaned and disinfected using an alcohol swab. Transposon plasmid (PT4-CMV-GFP (Addgene #11704), PT4-U6-sgRNA-CMV-GFP) and transposase overexpression plasmid pCMV(CAT)T7-SB100 (Addgene #34879) was allowed to equilibrate at room temperature (RT), transferred to a 3 mL syringe mounted with a 27G needle, and then injected into the tail vein in a continuous motion. The mouse was removed from the restrainer and signs of recovery were monitored.

Intrasplenic injection of tumor cells followed by splenectomy: Intrasplenic injection of tumor cells followed by splenectomy was previously described^88^ Briefly, mice were placed in a container connected to an inhalation-oxygen/isoflurane-device (4% for induction and 2-3% for maintenance). After 3-5 min, anesthetized mice were placed on a 37 °C thermal pad, isoflurane gas was continuously supplied by a nose cone, and sterile eye ointment was applied to avoid corneal dehydration. Anesthesia depth was monitored regularly by testing toe and tail pinch reflexes and by observing rate, depth and pattern of respiration. All surgical instruments were sterilized prior to use and the surgical procedure was performed under aseptic conditions. The skin over the surgical site was shaved and disinfected with Betadine. Using aseptic technique, an incision was made in the skin and peritoneal wall to expose the spleen. A sterile gauze was placed under the spleen. For each mouse, four 50 μL domes of AKPS organoids were harvested, washed from Matrigel in ice-cold PBS and mechanically dissociated using a 20 G needle on a 20 mL syringe. Dissociated organoids were pelleted at 290 g for 3 min, and resuspended in 0.04 mL sterile PBS and injected under the splenic capsule with an insulin syringe (BD, MicroFine, 0.3 mL, 30 G). Alternatively, 100’000 KPC cells in 0.04 mL sterile PBS were injected. After 10 minutes, the splenic artery and vein were closed by ligation. Immediately thereafter, the spleen was resected. Subsequently, the wound was washed 3 times with sterile PBS. The peritoneal wall was closed with an absorbable polyglactin suture (Vicryl 4-0 or 5-0 coated) and the skin was closed with wound clips. Mice were monitored for weight loss and the experiment was terminated maximally 3 weeks after tumor cell injection.

Tail vein injection of adeno-associated virus 8 (AAV8): A 100 μL solution of 1×10^11^ - 2×10^11^ AAV viral genomes in sterile saline was loaded into a 1 mL syringe mounted with a 27G needle. Mice were placed in a restraining device, and the tail was warmed with a red light lamp to induce lateral tail vein dilatation. The injection site was cleaned and disinfected using an alcohol swab, and then the AAV-containing solution was injected into the tail vein.

### In vivo CRISPR-a screen

Library cloning: sgRNA sequences for dCas9-SPH-mediated overexpression were retrieved from the Caprano library^89^ and obtained as oPool from Integrated DNA Technologies with Gecko flanking sequences^90^. Libraries were cloned into the transposon vector PT4-CMV-GFP (Addgene #117046) and amplified as described in^90^. Briefly, oPools were resuspended to 1 μg/μL in water and incubated for 2 h at 37°C, then amplified by polymerase chain reaction (PCR, See Supplementary Table S2 for primer sequences). Briefly, 1 uL library (1 ng/uL) was mixed with 12.5 uL NebNext MasterMix, 1.25 uL Oligo reverse primer (10 uM), 1.25 uL Oligo forward primer (10 uM) and 9 uL water and incubated in a thermocycler with the following program: 98 °C for 30s, 98 °C for 10 s, 63 °C for 10 s, 72 °C for 12 s, back to step 2 for a total of 20 cycles, 72 °C for 2 min. The PCR products were purified with Qiagen QIAquick PCR Purification Kit, eluted in 30 μL buffer EB (Qiagen) and separated on a 2% agarose gel in Tris-borate EDTA (TBE) buffer with SybrSafe Dye. The transposon vector was digested overnight with the Bsmb-v2 enzyme and run on a 2% agarose gel in TBE buffer with SybrSafe Dye. The 150 bp sgRNA amplicon and the digested vector band (missing 1000bp filler) were excised and gel-extracted using QIAquick Gel Extraction Kit. Both vector and insert were subjected to isopropanol purification by incubating 50 μL eluted DNA with 50 μL isopropanol, 0.5 μL GlycoBlue Coprecipitant and 0.4 μL 5M NaCl. Reactions were vortexed and incubated at room temperature (RT) for 15 min, prior to centrifugation at 16000 g for 15 min at RT. The precipitate was washed twice with 1 mL ice-cold 80% EtOH and air-dried for 1 min before resuspension in 10 μL water. DNA concentration was measured at Nanodrop. The Gibson assembly reaction mix containing 10 μL MasterMix, 330 ng vector, 50 ng insert, and water to 20 μL was incubated at 50 °C for 1 h. Following a second isopropanol precipitation, the cloned transposon libraries were resuspended in 5 μL Tris-EDTA (TE) buffer and incubated at 55 °C for 10 min. DNA concentration was measured at Nanodrop.

Library amplification: E.cloni electrocompetent bacteria (Biosearch Technologies) were thawed on ice for 20 min prior to addition of 1 μL of Gibson reaction and electroporation (1800 V) in a MicroPulser Electroporator (Biorad). 975 μL of pre-warmed recovery medium were added and the bacteria were incubated 37 °C at 250 rpm for 1 hour. Bacteria were plated on pre-warmed 30cm^2^ square agar plates with ampicillin and incubated overnight at 37 °C. Plates were scraped with 30-50 mL Luria-Bertani (LB) broth and plasmids were purified with Endotoxin-Free MaxiPrep kit (Macherey-Nagel). Precipitated DNA was resuspended in 1 mL water and the concentration was measured at Nanodrop. pCMV(CAT)T7-SB100 (Addgene #34879) was amplified in competent E. coli under chloramphenicol selection and purified with the Endotoxin-Free MaxiPrep kit (Macherey-Nagel).

Library delivery to mouse liver and tumor inoculation: the sgRNA transposon plasmid libraries (150 μg) and SB100X-encoding plasmids (15 μg) were co-injected hydrodynamically into AlbCre;dCas9-SPH mice or littermate controls lacking AlbCre. One week later, dissociated AKPS^sLP-mCherry^ organoids were inoculated by intrasplenic injection, followed by splenectomy. Metastases were allowed to grow for 2 weeks.

Isolation of metastasis-distal and proximal hepatocytes: mice were sacrificed by raising CO_2_ concentrations, then the abdomen was opened and a G22 cannula was inserted into the inferior vena cava. The liver was perfused with 20 mL Hanks buffer (0.5 mM EDTA and 25 mM HEPES in HBSS) followed by 15 mL digestion buffer (15 mM HEPES and 32 μg/mL Liberase in low glucose DMEM). After initial swelling of the liver, the portal vein was cut to allow outflow. After perfusion, the gallbladder was removed and the liver was transferred to a petri dish with 10 mL digestion buffer and squished with a cell scraper to release the hepatocytes. Liberase was inactivated by adding 10 mL isolation buffer (10% fetal bovine serum (FBS) in low glucose DMEM). The cell suspension was filtered through a 100 μm cell strainer and centrifuged at 50 g for 2 min. The supernatant was removed and the pellet was washed again twice with 20 mL isolation buffer. Hepatocytes were resuspended in 2 mL FACS buffer (2 mM EDTA, 0.5% BSA in PBS) and stained with Zombie Violet (1:500) (Biolegend #423113), TruStain FcX™ (anti-mouse CD16/32) antibody (BioLegend #101320, 1:50), PE/Cy7 anti-mouse CD31 (BioLegend #102418, 1:300), BV570 anti-mouse CD45 (BioLegend #103135, 1:300) for 25 min at 4 °C. Hepatocytes were washed and filtered through a 70 μm strainer. CD45^-^ CD31^-^ hepatocytes which contained a sgRNA (GFP^+^) were divided into metastasis-proximal (mCherry^+^) and metastasis-distal (mCherry^-^) by drawing different sorting gates on an AriaIII sorter (BD Biosciences) with 70-micron nozzle. Cells were collected in PBS, spun down at 800 g for 5 min and the pellet was stored at -20 °C.

Genomic DNA extraction and targeted guide amplification and sequencing. Genomic DNA extraction was performed as described in the “Isolation of genomic DNA with NucleoSpin Blood Kits and PCR pre-check” protocol of the Broad Institute’s Genetic Perturbation Platform. Briefly, the pellet was equilibrated at RT, resuspended in 200 μL PBS and incubated with proteinase K and lysis buffer mixture at 70°C overnight. 1 μL RNAse A was then added for 5 min at RT, followed by column purification with the NucleoSpin Blood Mini kit (Macherey Nagel). DNA was eluted in 25 μL elution buffer (prewarmed at 70 °C), incubating on column for 5 min prior to centrifugation, then the concentration was measured at Nanodrop. sgRNAs were target amplified from both post-injection libraries and plasmid library using an equimolar mixture of staggered P5 primers and P7 primers with sample-specific barcode (sequences in Supplementary Table S2) under the following conditions: 10 μL Titanium Buffer, 8 μL dNTPs, 5 μL DMSO, 0.5 μL P5 primer mix (100 μM), 40 μL water, 1.5 μL Titanium Taq polymerase, 25 μL DNA (max 10 ug), 10 μL P7 primer (5 uM). The reactions were incubated in a thermocycler with the following program: 95 °C for 5 min, 95 °C for 30 s, 53 °C for 30 s, 72 °C for 20 s, back to step 2 for a total of 28 cycles, 72 °C for 10 min. PCR products from different samples were pooled according to the number of reads required to ensure 200-1000 reads per sorted cell. After 1X SPRIselect bead (Beckman Coulter #B23319) purification, PCR products were eluted in 50 μL TE buffer. Quality and quantity libraries were assessed using dsDNA high-sensitivity kit (Life Technologies #Q32854) on a Qubit 4 fluorometer (Thermo Fisher) and the high sensitivity D1000 reagents and tapes (Agilent #5067-5585, #5067-5584) on a TapeStation 4200 or Bioanalyzer (Agilent Technologies). Libraries were sequenced on a NextSeq kit (Illumina) with 75bp single-end read chemistry and 9 bp index read, adding 10% PhiX spike-in (Illumina).

Replicates and coverage: the procedure outlined above (from cloning to sequencing) was repeated independently 3 times (3 batches). In sum, the sgRNA coverage (i.e. the number of sorted cells / 997 sgRNAs) added up to a total of 1000X for AlbCre;dCas9-SPH mice (n = 7) and 500X for nonCre littermate controls (n = 5).

Analysis: FASTQ files were demultiplexed with Bcl2fastq v2.20.0.422 (Illumina) and adaptors were trimmed with cutadapt^91^ with following parameters (-g CACCG and -a GTTTT). Sequences were aligned to the sgRNA library with Bowtie2^92,93^. sgRNAs were counted with the MAGeCK count function (--norm-method total)^94,95^. sgRNA enrichment was calculated with the MAGeCK paired test function. Metastasis-proximal and metastasis-distal libraries from the same mouse were considered as paired samples. As sgRNAs in paired samples are considered as independent sgRNAs (3 sgRNA in 7 mice are considered 21 independent sgRNAs), paired testing allows to find consistent effects between paired replicates. This analysis was repeated for the individual library batches. The paired function was not used to compare perturbation enrichment in AlbCre;dCas9-SPH vs noCre littermates, as an equal number of samples is required. Instead, sgRNAs counts were added up from all mice and then the standard MAGeCK RRA test function was applied. Results were visualized with MAGeCKFlute^94^ and ggplot2^96^.

### Cross-validation with human mutational data

Interactors of liver metastases-specific mutations: The Genomic features of organotropism dataset^39^ was used to extract the mutations associated with liver metastases. This set of genes was then parsed with CellPhoneDB v3^40^, CellTalkDB^41^, and NicheNet^38^ to screen for ligand and receptors and compile a list of all their known interactors. We then filtered for genes expressed by hepatocytes according to the Human Protein Atlas (v22.0 proteinatlas.org)^97^). Specifically, we filtered out genes with transcript per million (TPM) μ 0.5 in the RNA GTEx tissue gene data, normalized TPM (nTPM) μ 0.5 in the RNA single cell type data (v22.0 https://www.proteinatlas.org/humanproteome/single+cell+type), and “not detected” in the Normal tissue data (v22.0 proteinatlas.org, Ensembl version 103.38). The data used for the analyses described in this manuscript were obtained from the GTEx Portal on 05/30/2022 and/or dbGaP accession number phs000424.vN.pN on 05/30/2022.

Kaplan-Meier analysis: Kaplan-Meier analysis of 1211 CRC patients was performed using the online tool http://kmplot.com. Recurrence free survival (RFS) was stratified by SEMA4A, SEMA4C and SEMA4D expression in the Affymetrix colon dataset, using best cutoff.

Visium library preparation: A sample of human CRC hepatic metastasis (#CB522586, 44 years old male) with clear tumor-liver borders was selected from a commercial biobank (Origene). Non-consecutive sections were cut with a thickness of 10μm and placed onto 2 capture areas of 10X Visium Spatial Gene expression slide using a cryostat (Leica CM3050S). The tissue Optimization kit was used to determine permeabilization times (24 min), and cDNA libraries were generated following manufacturer’s instructions (10x Genomics). Quality and quantity of libraries were assessed using dsDNA high-sensitivity kit (Life Technologies #Q32854) on a Qubit 4 fluorometer (Thermo Fisher) and the high sensitivity D1000 reagents and tapes (Agilent #5067-5585, #5067-5584) on a TapeStation 4200 (Agilent Technologies). Paired-end sequencing was performed for all libraries (read 1:28 bp, index read: 10 bp, read2: 82 bp) on a NovaSeq 6000 (Illumina) using NovaSeq SP Reagent Kits (100 cycles).

Data analysis: Binary base call (BCL) files were demultiplexed using Bcl2fastq v2.20.0.422 (Illumina) and pre-processed using Space Ranger (v1.1.0 or v1.2.0, 10x Genomics). Spot transcriptomes were deconvoluted with Spotlight^98^ using published single-cell RNA sequencing data as reference. Specifically, we integrated two datasets of primary colorectal cancer^36^ and liver tumor microenvironment^37^ using the Seurat integration method^99^. Edge spot selection was performed with the CellSelector function in Seurat and the FindMarkers function was used for differential gene expression analysis of metastasis-proximal vs distal and metastasis-core vs center. NicheNet^38^ was used to predict ligand-receptor interactions at the metastatic leading edge. The ligand activity analysis from NicheNet was used to estimate the potential of these interactions to regulate the observed transcriptomic alterations.

### Spatial transcriptomics of human CRC hepatic metastases

Mouse CRC organoid culture: VilCreER^T2^;APC^fl/fl^;Tp53^fl/fl^; Kras^G12D/wt^Smad4^KO^ (AKPS) organoids were cultured in 50 domes μL of Matrigel (Corning) as described previously^101^. To make complete medium, Advanced DMEM/F12 (Life Technologies™) was supplemented with 10 mM HEPES (Life Technologies™), 2 mM L-Glutamine (Life Technologies™), 100 mg/mL Penicillin/streptomycin (1% PenStrep), 1x B27 supplement (Life Technologies™), 1 x N2 supplement (Life Technologies™) and 1mM N-acetylcysteine (Sigma-Aldrich). Organoids were split every 3-5 days by mechanical dissociation. Splitting was always performed on the day preceeding intrasplenic injection.

RNP-mediated Smad4-KO: VilCreER^T2^;APC^fl/fl^;Tp53^fl/fl^;Kras^G12D/wt^ were obtained from Owen Sansom (Beatson Institute for Cancer Research, Glasgow, UK) and cultured under the above conditions with supplementation of 100 ng/mL murine recombinant Noggin (LuBioScience, #250-38-250). 4 domes of organoids were treated with 5 mM nicotinamide for 2-3 days prior to the experiment. sgRNAs comprising both crRNA and tracrRNA sequences were obtained from IDT (Alt-R system). Targeting sequences were obtained from^102^. Organoids were harvested, washed from Matrigel, and dissociated into single cells by incubation at 37 °C for 10 minutes in 1 mL pre-warmed TrypLE™ Express Enzyme (Gibco). Cells were spun down at 190 g for 3 min, resuspended in 1 mL complete medium with 5 mM nicotinamide and 10 μM Y-27632 dihydrochloride (Rock inhibitor) (Stemcell #72304) and seeded in 2 wells of a 48-well plate. Transfection was performed with the CRISPRmax kit (Thermo Fisher Scientific). Briefly, 25 μL OPTImem supplemented with 1250 ng Cas9 nuclease V3 (#1081058), 240 ng sgRNA and 2.5 μL Cas9 Reagent Plus were mixed with 25 mL OPTImem with 1.5 μL CRISPRmax reagent, incubated for 10 min and added to each well. The cells were spinoculated for 1 hr at 600 g at 32 °C, then incubated 4 hrs at 37 °C. Cells were then harvested, resuspended in Matrigel and plated in complete medium supplemented with Rock inhibitor. 3 days later, selection was started by adding medium supplemented with 10 ng/mL TGF-β (Peprotech #100-21C-10UG) and lacking Noggin. Selection was continued for 3 passages, then TGF-β was removed from the medium. Successful editing was confirmed by T7 endonuclease assay. Briefly, primers asymmetrically flanking the cut site were designed so as to yield fragments distinguishable by electrophoresis. Organoid DNA was extracted with QuickExtract™ DNA Extraction Solution (Lucigen) and PCR amplified. The PCR reaction mix consisted of 10 μL Q5 Master mix, 6 uL H_2_O, 2 μL primer mix and μL DNA. 10 uL PCR products were mixed with 1.5 μL 10x NEB Buffer 2 and 1.5 μL H_2_O and incubated in a thermocycler as follows: 10min 95 °C, from 95 °C to 85 °C with ramp rate of -2°C/sec, from 85 °C to 25 °C with ramp rate of -.3C/sec. Formed heteroduplexes were incubated with 2 μL T7 mix (10 μL NEBuffer 2, 10 μL T7 and 80 μL H2O) at 37 °C for 1h. The reaction was stopped by the addition of 1μL of 0.5 M EDTA. Samples were analyzed by electrophoresis on a 2% agarose gel. Abrogation of SMAD4 signaling was further confirmed loss of Id3 expression, as assessed by qRT-PCR.

Integration of sLP-mCherry labeling system: the pcPPT-mPGK-attR-sLPmCherry-WPRE vector was obtained from ximbio.com and lentiviruses were generated according to a published protocol^103^. Briefly, HEK-293T cells were cultured on poly-D-lysine coated plates and transfected with 4.4 μg PAX2, 1.5 μg VSV-G and 5.9 μg lentiviral vector with JetOptimus reagents. The supernatant was harvested after 2 days, spun down 5 min at 500 g and filtered through a 0.45 μm filter. 4 domes of AKPS organoids were washed from Matrigel and dissociated into single cells by incubating them for 10 minutes in 1 mL pre-warmed TripLE at 37 °C. Cells were centrifuged at 190 g for 3 min, then resuspend in 2 mL infection medium containing 1.8 mL virus, 200 μL complete medium, 5 mM nicotinamide, 1.6 μL polybrene and 2 μL Rock inhibitor. The cells were plated in 2 wells of a 48 well plate and spinoculated for 1 hr at 600 g at 32 °C, then incubated 4 hrs at 37 °C. Cells were then harvested, resuspended in Matrigel and plated in complete medium supplemented with Rock inhibitor. 3 days later, organoids were dissociated into single cells and mCherry^+^ cells were isolated by FACS using an AriaIII sorter (BD Biosciences). Organoids were further selected with 2μg/mL puromycin for a week.

shRNA-mediated semaphorin KD: sequences of shRNAs targeting Sema4a, Sema4c, Sema4d and Sema4g were obtained from the The RNAi Consortium shRNA Library (Broad Institute, Cambridge, MA, United States) and cloned in an arrayed fashion in a lentiviral vector expressing GFP as a selection marker based on a published plasmid backbone^104^. The empty vector control expressed a puromycin-resistance cassette for selection. Lentiviral preparation and transduction of organoids was conducted as above. Organoids transduced with shRNAs targeting semaphorins (Sema-KD) were dissociated into single cells, incubated with Zombie Violet (1:500, Biolegend #423113) and selected by FACS based on GFP fluorescence and gating for live cells. AKPS organoids transduced with the empty vector were also dissociated into single cells and subjected to sorting (only live cell gate), and then selected by puromycin as described above. Semaphorin knockdown was assessed by qRT-PCR. For competitive seeding assays, Sema-KD and empty vector organoids were grown separately and then mixed in a 1:1 ratio and mechanically dissociated for intrasplenic injection. A small fraction of the injection mix was seeded in 3 domes to estimate injection ratios.

Patient-derived CRC organoids: human CRC organoids were obtained from the Visceral Surgery Research Laboratory at the University of Basel. CRC-patient tissues from primary and liver metastases were obtained from the University Hospital Basel following patient consent and ethical approval (Ethics Committee of Basel, EKBB, number EKBB numbers 2019-00816). Patient-derived organoids were generated as previously described (PMID: 33754045). Briefly, tissue was cut into small pieces and, subsequently, enzymatically digested in 5 mL advanced DMEM/F-12 (ThermoFisher Scientific, #12634028) containing 2.5 mg/mL collagenase IV (Worthington, #LS004189), 0.1 mg/mL DNase IV (Sigma, #D5025), 20 ug/mL hyaluronidase V (Sigma, #H6254), 1% BSA (Sigma, #A3059) and 10 μM LY27632 (Abmole Bioscience, #M1817) for 1 hour and 30 minutes at 37°C under slow rotation and vigorous pipetting every 15 minutes. The tissue lysate was filtered through a 100 μM cell strainer and centrifuged at 300 g for 10 minutes the cell pellet was finally resuspended with growth factor reduced Matrigel (Corning, #356231) and plated into 20 μl domes. After polymerizationl, Matrigel embedded cells were overlaid with medium composed of advanced DMEM/F-12 supplemented with 10 mM HEPES (ThermoFisher Scientific, #15630056), 100 μg/ml Penicillin-Streptomycin (ThermoFisher Scientific, #10378-016), 1” Glutamax (ThermoFisher Scientific, #9149793), 100 μg/ml primocin (invivoGen, #antpm-1), 1x B27 (ThermoFisher Scientific, #17504044), 1.25 mM N-acetyl-cysteine (Sigma-Aldrich, #A9165-25G), 10 mM Nicotinamide (Sigma-Aldrich, #N0636), 500 ng/ml R-Spondin (EPFL SV PTPSP), 100 ng/ml Noggin ((EPFL SV PTPSP), 50 ng/ml EGF (PeproTech, #AF-100-15), 500 nM A83-01 (R&D Systems, #2939), 10 μM SB202190 (Sigma-Aldrich, #S7076), 10 nM prostaglandin E2 (Tocris Bioscience, # 2296), 10 nM gastrin (Sigma-Aldrich, # G9145) and 10 μM Y-27632 dihydrochloride. Medium was changed every 3 days, and organoids were passaged using 0.25% Trypsin-EDTA ((Life Technologies, #25200-056) or Gentle Cell Dissociation Reagent (StemCell, #100-0485).

### Cell lines

Mouse immortalized hepatocytes: AML12 cells were obtained from ATCC (#CRL-2254) and cultured in DMEM:F12 Medium (Gibco) supplemented with 10% FBS, 10 μg/ml insulin, 5.5 μg/ml transferrin, 5 ng/ml selenium, 40 ng/ml dexamethasone and 1% PenStrep. dCas9-SPH was introduced by recombinase-mediated cassette exchange (RCME)^105^ in the Rosa26 safe-harbor^106^.

Human immortalized hepatocytes: PTA-5565 cells (ATCC) stably labeled by H2B-mCherry were obtained from the Bentires-Alj laboratory (University of Basel) and cultured in William’s E Medium supplemented with 1% Glutamax (Gibco), 5% FBS and 1% PenStrep.

KPC cells: Ptf1a-Cre;Kras^G12D/+^;Trp53^flox/+^ (KPC) pancreatic ductal adenocarcinoma cells (B6J syngeneic) were kindly donated by Ilaria Guccini (ETH Zurich) and cultured in DMEM:F12 supplemented with 10% FBS and 1% PenStrep. Cells were split every 3-5 days and on the day before surgery. Before intrasplenic injection, KPC cells were detached from culture flasks with 1mM EDTA.

### In vitro arrayed screen and seeding assays

Arrayed screen with primary hepatocytes or AML12 cells: 96 or 384 well plates were coated with rat tail collagen with the Collagen-I Cell Culture Surface Coating Kit (ScienCell Research Laboratories) following manufacturer’s instructions. Primary mouse hepatocytes from AlbCre;dCas9-SPH mice were isolated by perfusion as described above. After 2 washes in isolation buffer, the hepatocyte pellet was further purified by density separation following a published protocol^107^. Briefly, the pellet was resuspended in 10 mL isolation buffer and 10 mL Percoll solution (9 mL Percoll, 1 mL 10X PBS), then mixed thoroughly by inverting the tube several times. Following centrifugation at 200 g for 10 min at 4°C, the hepatocytes were resuspended in isolation medium (supplemented with 1% PenStrep) and plated at high density (15’000 hepatocytes per well in 96 well plates, 5’000 hepatocytes per well in 384 well plates). The same plating density was used for AML12 cells expressing dCas9-SPH. On the next day, primary or immortalized hepatocytes were transfected with SB100X and a pool of 3 transposon vectors harboring sgRNAs against selected gene targets using Lipofectamine 3000 (Thermo Fisher Scientific). Three wells were independently transfected for each target in an arrayed fashion, and three wells were left untransfected. On the next day, transfection efficiency was estimated by GFP fluorescence. AKPS^sLP-mCherry^ organoids were dissociated into single cells as described above, then 50 cells were seeded per well. After 5 days, colony formation was assessed by microscopy.

Stable overexpression of Plxnb2: AML12-cells expressing dCas9-SPH were transfected with SB100X and a pool of 3 PT4-U6-sgRNA-CMV-GFP transposon vectors targeting Plxnb2, or sgNT, using Lipofectamine 3000 (Thermo Fisher Scientific). On the next day, cells were sorted for GFP^+^ fluorescence on AriaIII sorter (BD Biosciences) with 70-micron nozzle and used for colony formation assays as described above.

Treatment with recombinant mouse and human Plexin B2: Recombinant human Plexin B2 (5329-PB-050, Biotechne) and mouse Plexin B2 (6836-PB-050, Biotechne) were reconstituted at 100 μg/mL in PBS. Human or mouse organoids were dissociated into single cells as desribed above, mixed with recombinant Plexin B2 in growth medium, then seeded in 384 well plates at a density of 50 cells/well, in the absence or presence of 5000 human or mouse hepatocytes. Colony formation was scored by microscopy. Were indicated, cultures were supplemented with 50 uM Rac1 inhibitor NSC 23766, 1 ng/uL anti-SEMA4D monoclonal antibody (Pepinemab Biosimilar, Proteogenix PX-TA1382 or 1 ng/uL anti-PLXNB2 monoclonal antibody (67265-1, Proteogenic). The EdU incorporation assay was performed with the Click-iT EdU Cell Proliferation Kit for Imaging, Alexa Fluor 647 dye (Invitrogen). Briefly, 3 days after treatment of PDOs with rhPlexin B2, half of the culture medium was removed and replaced with 2X EdU-containing medium for 1 hr, then the manufacturer’s instructions were followed.

### Adeno-associated virus generation

Three sgRNA sequences for dCas9-SPH-mediated overexpression (OE) of Plxnb2 were obtained from the Caprano library^89^, four sgRNA sequences for Cas9-mediated knockout (KO) and two control non-targeting sgRNA were obtained from the Brie library^108^. See Supplementary Tables S1 and S2 for sequences. Each guide was cloned into U6-sgRNA-EF1a-EGFP (Addgene #117046) vector and amplified using the Maxi prep kit (Macherey-Nagel). 250 million HEK-293T cells were seeded in a Five Chambers Cell-Stack (Corning). 24 hours later, sgRNA vectors for each given condition (OE, KO, NT) were pooled and co-transfected with pAdH helper plasmid and pAAV2/8 capsid (Addgene, #112864) with polyethylenimine (PEI). Adeno-associated viruses (AAVs) were generated and purified following a slightly modified version of the AddGene protocols “AAV Production in HEK293T Cells” and “AAV Purification by Iodixanol Gradient Ultracentrifugation”. Following viral genome production and purification, total viral genomes were quantified using digital droplet PCR (ddPCR) according to the Addgene protocol “ddPCR Titration of AAV Vectors”. AAVs were injected in the tail vein as described above, 1 week prior to AKPS organoids or KPC cells intrasplenic injections.

### Immunofluorescence

Formalin-fixed and embedded tissues: Livers were perfused with PBS, then the medial lobe was fixed with 4% PFA in PBS for 48 hours, followed by 48 hours PBS incubation and storage in 75% EtOH. Dehydration, formalin embedding and H&E staining were performed by the histology core facility of the University of Basel. 5 μm sections were deparaffinized with descending alcohol series and subjected to heat-induced epitope retrieval in 2.4 mM sodium citrate and 1.6 mM citric acid, pH 6, for 25 min in a steamer. Sections were washed with PBST (0.1% Tween-20 in PBS) and blocked for 1 h at RT in blocking buffer (5% BSA, 5% heat-inactivated normal goat serum in PBST). Sections were incubated overnight at 4 °C with the following primary antibodies (1:100, in blocking buffer): anti-CD146 (Abcam #ab75769); anti-#-SMA (Abcam #ab5694) anti-periostin (Abcam #ab227049) and anti-GFP (Aves Labs #GFP-1020). Sections were repeatedly washed in PBST and incubated with the following secondary antibodies (1:400, in blocking buffer) for 1 h at RT: AlexaFluor goat anti-rabbit 594 (#A-11012), AlexaFluor goat anti-rabbit 647 (#A-21244), AlexaFluor goat anti-chicken 647(#A32933), all from ThermoFisher. Nuclei were stained with DAPI (Sigma, 1:1000) in blocking buffer for 15 min at PBST. Sections were mounted with Prolog Gold (P36930 Invitrogen) and imaged on the Leica THUNDER Imager 3D Cell Imaging system, equipped with a Leica LED8 Light engine, Leica DFC9000 GTC sCMOS camera and the following filter sets: filter cube CYR71010 (excitation: 436/28, 506/21, 578/24, 730/40, dichroic: 459, 523, 598, 763, emission: 473/22, 539/24, 641/78, 810/80); filter cube DFT51010 (excitation: 391/32, 479/33, 554/24, 638/31, dichroic: 415, 500, 572, 660, emission: 435/30, 519/25, 594/32, 695/58), and extra emission filters (460/80, 535/70, 590/50, 642/80, 100%).

Fixed frozen tissue: Following liver perfusion with PBS, the left lobe was incubated in 4% PFA at 4 °C for 1 hour, and then in 30% sucrose in PBS overnight at 4 °C and then embedded in Tissue-Tek OCT Compound (Sakura, 4583) for cryosectioning. 8 μm sections were washed 3 times, blocked and stained as above with the following primary and secondary antibodies: anti-glutamine synthetase (1:100m, Biolegend #856201), anti c-Myc (9E10, Thermo Fisher Scientific), anti-LDLR (Biotechne, #MAB2255), anti-Plexin B2-PE (1:500, Biolegend #145903), anti-GFP-AlexaFluor488 (1:200, ThermoFisher #A-21311), anti-Zeb1 (1:400, Novus #NBP1-05987), anti-#-SMA (1:1000, SIGMA #A2547), DAPI counterstain, mounting and imaging was performed as above. F-actin was stained by incubating blocked slides for 2 hours at RT with Alexa Fluor 647 Phalloidin (1:400, Invitrogen #A22287).

Organoids: Following fixation in 4% PFA at 4 °C for 2 hours, and then blocked in 5% BSA-PBS solution with 0.2% Triton, followed by primary (overnight) and secondary antibodies (4 hours): anti-Zeb1 (1:400, Novus #NBP1-05987), anti-Ecadherin (1:200, BD Biosciences #610181), DAPI counterstain, mounted in 3% low melting agarose in glass-bottom plates followed by imaging.

### Quantification of metastatic foci and lesion area

H&E sections were imaged on the Leica DMi8 inverted microscope, equipped with a FLEXACAM C1 12 MP CMOS camera and analyzed using QuPath software^109^. Whole tissue area and single metastases were manually isolated, producing a measure for whole section and metastases area (μm^2^), and metastases number per section. Two sections were taken and analyzed per animal, and a mean was calculated for the number of metastasis foci per liver section.

### Single molecule in situ hybridization

Custom DNA smFISH probes for Plxnb2 were designed in house and synthesized by Biosearch Technologies containing a 3’ amine reactive group (see Supplementary Table S2 for list of probes). All probes were pooled and labeled with AlexaFluor 594 dye according to a previously published protocol^110^. Mouse tissues were collected and fixed with 4% PFA in PBS for 3 hours followed by overnight incubations in 30% sucrose, 4% PFA in PBS at 4°C. Fixed tissues were embedded in Tissue-Tek OCT Compound (Sakura, 4583). 8 μm tissue sections were sectioned onto poly-L-Lysine coated coverslips, allowed to adhere by drying at RT for 10 min, followed by 15 min fixation in 4% PFA and overnight permeabilization in 70% EtOH. Probe hybridization was performed according to a previously published protocol^110^. Images were acquired with a 63x oil immersion objective with NA=1.4 on the Leica THUNDER Imager 3D Cell Imaging system, equipped with a Leica LED8 Light engine, Leica DFC9000 GTC sCMOS camera. For quantification, 3-4x FOVs covering the entire width of the tissue were acquired for each sample and the images were processed using the Thunder deconvolution algorithm. Maximum intensity projections of the processed images were rendered using ImageJ. Dot counting to determine transcript numbers for each FOV was performed with FISHQuant^111^ using the automatic thresholding function and cell number was determined by segmenting and counting the nuclei using CellPose^112^. Spot counting and nuclei numbers were manually verified to ensure correctness. Thereafter the average number of spots per cell was measured by dividing the number of spots within the FOV by the number of nuclei.

### Quantitative real-time polymerase chain reaction (qRT-PCR)

RNA extraction from fresh organoids and liver tissue was performed with the Qiagen RNeasy purification kit. 1 ng of total RNA was reverse transcribed with the cDNA synthesis kit (Takara Bio Inc.) according to the manufacturer’s instructions. Expression of genes of interest was quantified with primers listed in Supplementary Table S2, by qRT-PCR using the Applied Biosystems SYBR Green Kit monitored by the QuantStudio3 system (Applied Biosystems). Samples were analyzed in technical triplicates and average cycle threshold values were normalized to Gapdh using the ΔΔCT method^113^.

### Bulk RNA-sequencing experiments and analysis

Library preparation: RNA was extracted from microdissected metastases or organoids as above, and libraries for bulk RNA sequencing were prepared using the mcSCRB-seq protocol^114^. Libraries were quality-controlled using dsDNA high-sensitivity kit (Life Technologies #Q32854) on a Qubit 4 fluorometer (Thermo Fisher) and the high sensitivity D1000 reagents and tapes (Agilent #5067-5585, #5067-5584) on a TapeStation 4200 (Agilent Technologies) and sequenced on a NovaSeq 6000 (Illumina) using NovaSeq SP Reagent Kits (100 cycles).

Analysis: Reads were demultiplexed with Bcl2fastq v2.20.0.422 (Illumina) and quality-checked with FastQC^115^. Adaptors were trimmed with cutadapt^91^. Data was processed using the zUMIs31 (v2.9.4) platform to convert reads to count matrices per sample. Differential gene expression analysis was performed with edgeR^116^. Gene set enrichment analysis on significantly differentially expressed genes (logFC > |2|, p < 0.01) using the Bioconductor package fgsea with default parameters^117^. Genes were ranked based on adjusted p-value taking into account directionality of the fold change with the formula ranking = $log10(P)/sign(log2ratio) (http://genomespot.blogspot.co.at/2016/04/how-to-generate-rank-file-from-gene.html). The Gene Ontogy and Hallmarks gene set collections from the Molecular Signatures Database were imported in R with the package msigdbr^118^.

### Statistical analysis

Statistical analysis and visualization were performed using R (Version 3.4.0, R Foundation for Statistical Computing Vienna, Austria), R Studio and Prism 8.2.0. Statistical significance tests were performed as described in each figure legend, and p values were adjusted for multiple testing.

## Data and Code availability

The sequencing data generated in this study are available at the Zenodo repository: 10.5281/zenodo.7737590. The code used in this study is available at https://github.com/CocoBorrelli/Mosaic-Liver.

---

**Extended Data Figure 1.**
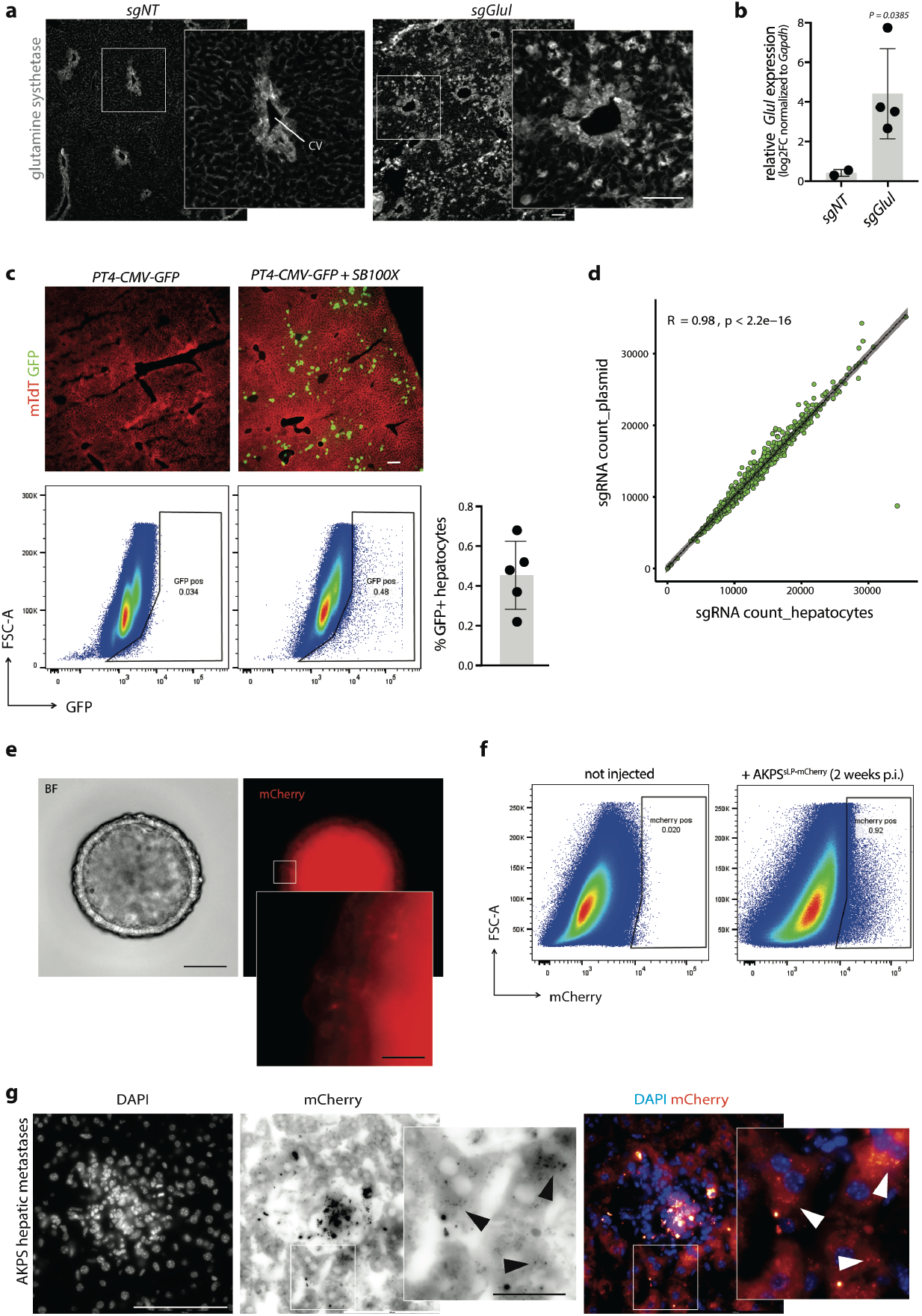
Screening tumor-hepatocytes interactions in a mosaic liver. **a**, Glutamine synthetase immunoreactivity in AlbCre;dCas9-SPH mice upon co-injection of *SB100X* and *PT4-U6-sgGlul* or *PT4-U6-sgNT*. Scale bar, 100 μm. CV, central vein. **b**, *Glul* gene expression of whole liver extracts, as assessed by qRT-PCR. Two-tailed unpaired t-test. n = 2. Log2FC normalized to Gapdh. **c**, Hydrodynamic tail vein injection (HTVI) of *SB100X* and *PT4-CMV-GFP* transposon. Top, endogenous GFP fluorescence. Scale bar, 100 μm. Bottom left, representative flow cytometry plot of GFP^+^ hepatocytes (gated as CD45^-^ CD31^-^) 1 week after co-injection of *SB100X* and *PT4-CMV-GFP*. Bottom right, percentage of GFP^+^ hepatocytes. *AlbCre;dCas9-SPH* mice (n = 5). Barplots indicate mean ± standard deviation (SD). **d**, sgRNA abundance in plasmid vs sorted GFP^+^ hepatocytes. Pearson’s correlation test. **e**, Representative micrograph of AKPS organoids expressing sLP-mCherry. Scale bar, 50 μm. Inset shows sLP-mCherry punctae. Scale bar, 10 μm. **f**, Representative flow cytometry plots of mCherry niche-labeling in vivo upon AKPS injection. Gated on CD31^-^CD45^-^ hepatocytes. **g**, Fluorescent micrograph of AKPS hepatic metastasis expressing sLP-mCherry 1 week post intrasplenic injection. Nuclei are labeled with DAPI. Scale bar, 100 μm. Arrowheads indicate sLP-mCherry punctae in labeled proximal hepatocytes. Scale bar, 20 μm.

**Extended Data Figure 2.**
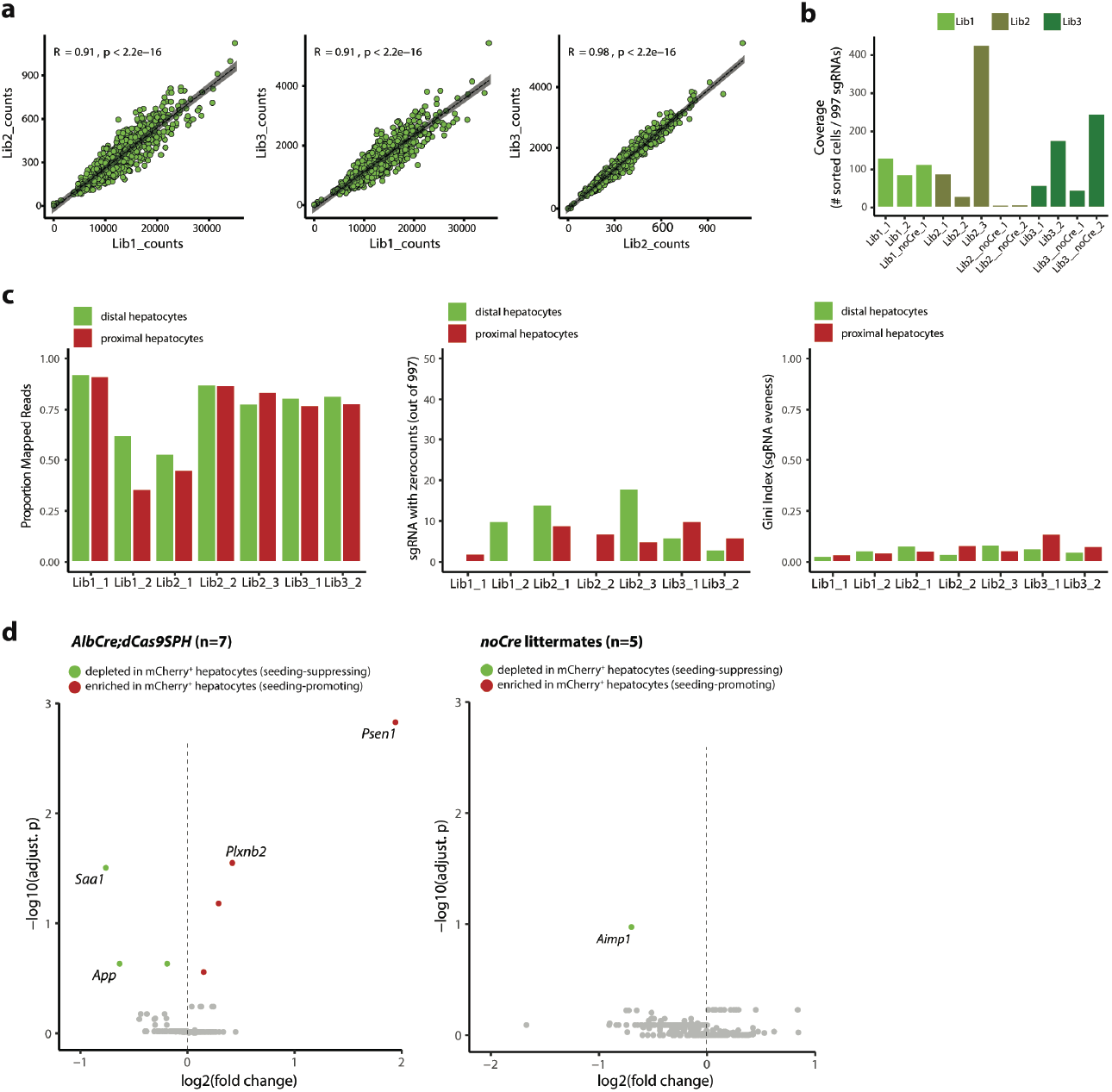
In vivo CRISPR-a screen identifies hepatocyte-derived factors that regulate metastatic seeding. **a**, Pearson correlation of sgRNA abundance in three independent library batches. **b**, sgRNA coverage per mouse, calculated by dividing the number of sorted cells by 997 (sgRNAs) (n = 7 *AlbCre;dCas9-SPH* mice and 5 *nonCre* littermate controls). **c**, Proportion of reads mapped (left), zero count sgRNAs (middle), and Gini index (right) of sgRNA libraries amplified from distal (mCherry^-^) and proximal (mCherry^+^) across three independent batches and recipient *AlbCre;dCas9-SPH* mice (n=7). **d**, Volcano plots showing screening results for *AlbCre;dCas9-SPH* mice (left) and nonCre littermate controls (right). Red dots indicate target genes enriched in the proximity of metastases (putative seeding-promoting factors). Green dots indicate target genes depleted in the proximity of metastases (putative seeding-suppressing factors).

**Extended Data Figure 3.**
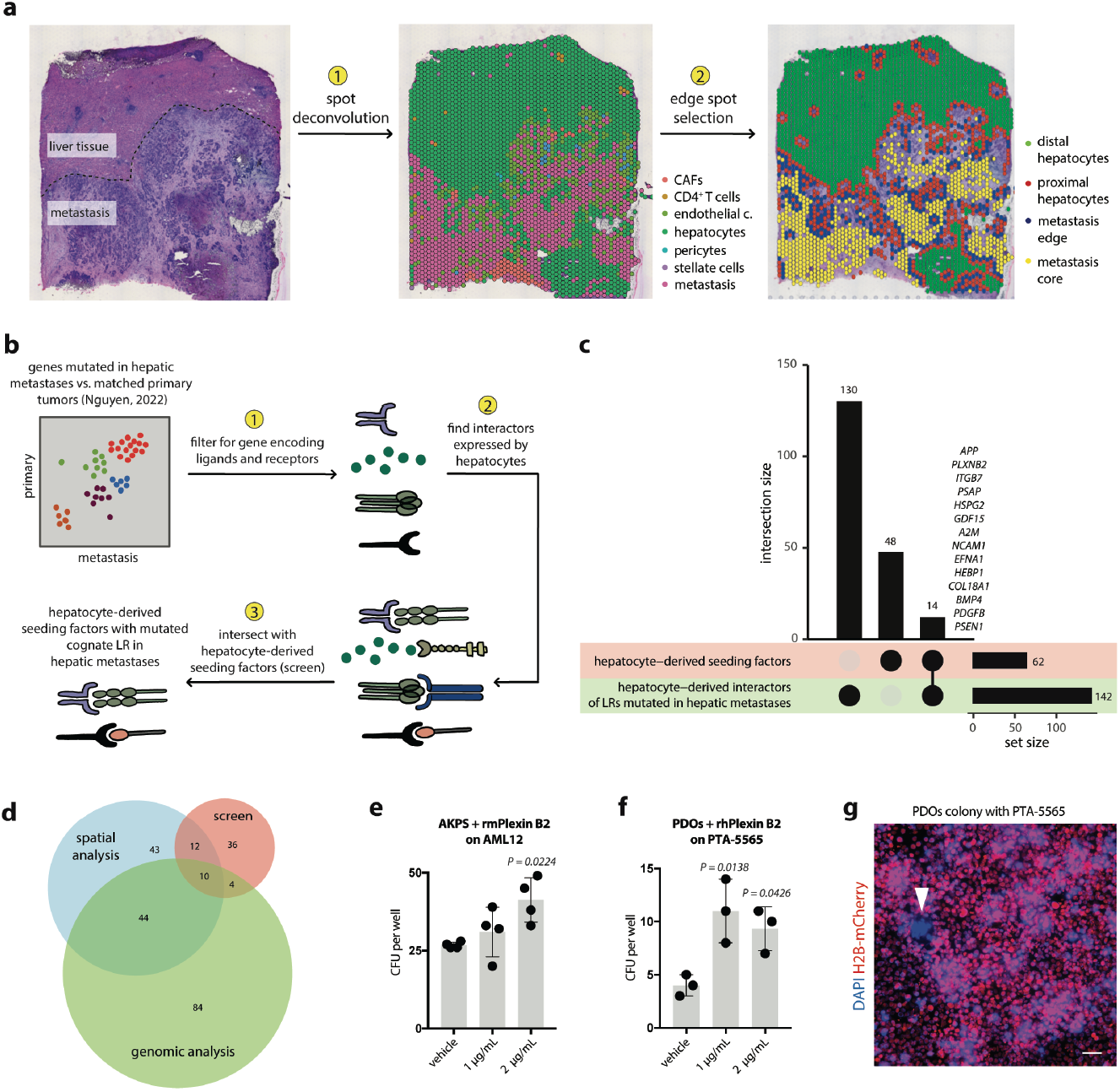
Cross-validation in human mutational and transcriptomic data. **a**, H&E staining (left) of CRC metastases from human liver used for Visium spatial transcriptomics (10x Genomics) (2 replicates). After 1) spot deconvolution, predominant cell types per spot are indicated in a color-coded map (middle). This is 2) used as input for edge spot selection, identifying metastasis-proximal and distal hepatocytes as well as metastasis core vs. edge. **b**, Schematic representation of analytical workflow. 1) Frequently mutated genes in hepatic metastases vs primary tumors in a publicly available dataset ^39^ are filtered for ligand or receptor-encoding genes. 2) Cognate interaction partners of identified genes are identified by cell-cell interaction prediction tools and filtered for hepatocyte expression. 3) Interaction partners of frequently mutated LRs in hepatic metastases are intersected with hepatocyte-derived seeding factors identified in the screen. **c**, Upset plot of intersection analysis. Intersecting genes are listed. **d**, Venn diagram shows total set and interaction sizes of hepatocyte-derived seeding factors identified in the screen (“screen”, red), predicted metastases-hepatocyte LR interactions at the tumor edge by spatial analysis (“spatial analysis”, blue), or cognate LRs of genes frequently mutated in metastases (“genomic analysis”, green). **e**, CFU per well of AKPS organoids seeded as single cells on AML12 cells in the absence or presence of increasing concentrations of rmPlexin B2. **f**, CFU per well of patient-derived CRC organoid (PDOs) cultured on PTA-5565 cells with increasing concentrations of rhPlexin B2. **g**, Representative fluorescent micrograph of a PDO colony (indicated by arrowhead) growing on H2B-mCherry labeled PTA-5565 cells. Scale bar, 20 μm. D-E, Barplots indicate mean ± SD. Dots represent individual wells. Ordinary one-way ANOVA, comparing each treatment group to vehicle control. Tukey’s correction for multiple testing.

**Extended Data Figure 4.**
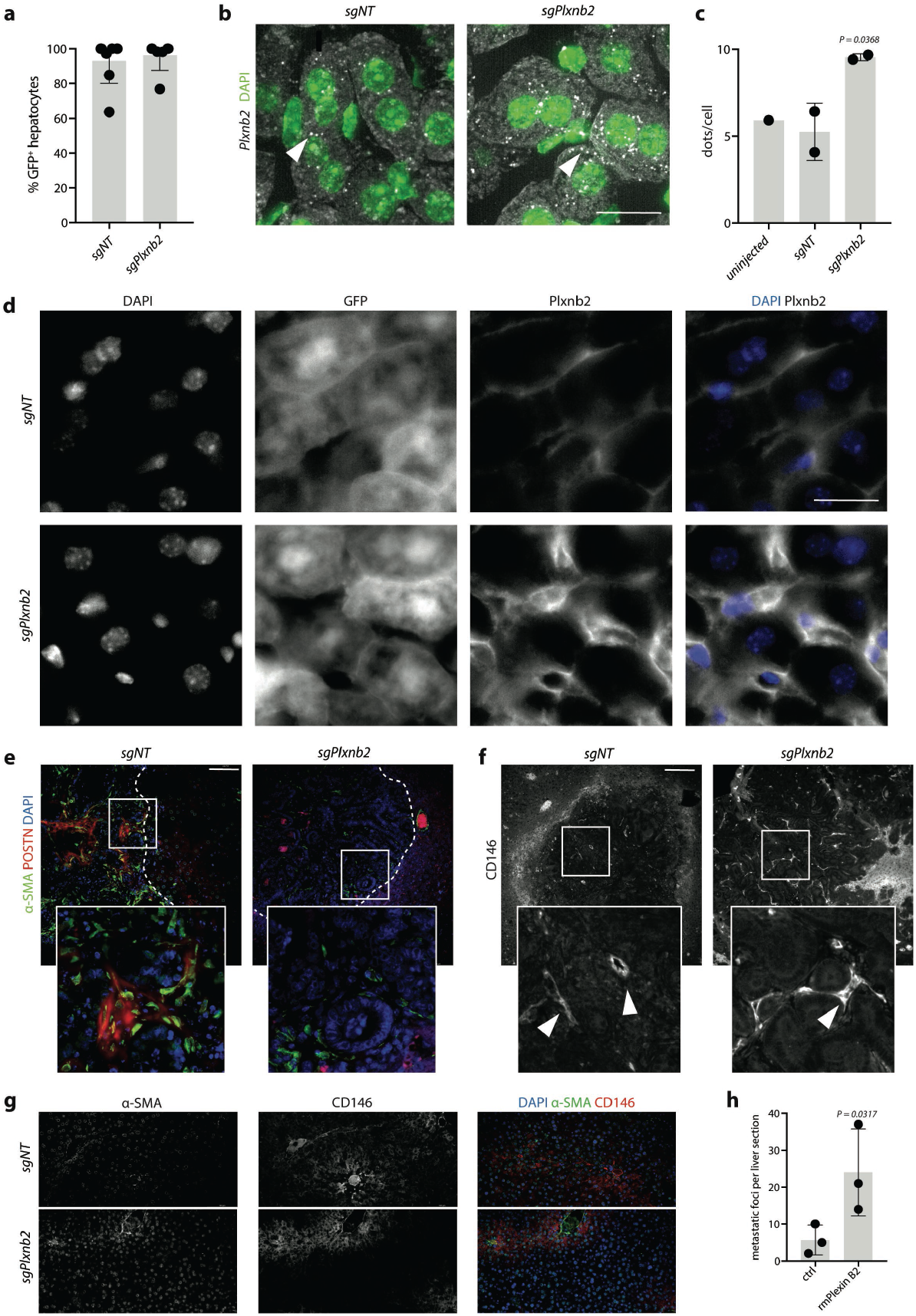
Overexpression of Plxnb2 in murine hepatocytes after AAV-mediated sgRNA transduction. **a**, Percentage of GFP^+^ hepatocytes as assessed by immunofluorescence in *AlbCre;dCas9-SPH* mice injected with *AAV8-U6-sgPlxnb2-EF1A-EGFP* (*sgPlnxb2*, n=5) or *AAV8-U6-sgNT-EF1A-EGFP* (*sgNT*, n=7). Scale bar, 20 μm. **b**, Representative micrograph of DAPI staining (green) and *Plxnb2* mRNA (white) as detected by smFISH. Arrowheads indicate single mRNA molecules. **c**, Quantification of Plxnb2 mRNA dots per cell in uninjected, *sgNT*- or *sgPlxnb2*-injected *AlbCre;dCas9-SPH* mice. **d**, Representative fluorescent micrographs of sgNT and sgPlxnb2 transfected livers showing nuclear DAPI staining, endogenous GFP signal and Plexin B2 immunoreactivity. Scale bar, 20 μm. **e**, Representative fluorescent micrographs of AKPS metastases in *sgPlxnb2* transduced livers showing nuclear DAPI stain (blue), and “-SMA (green) and periostin (POSTN, red) immunoreactivity. Dashed line indicates the tumor border. Scale bar, 20 μm. **f**, Immunofluorescence staining for the pericyte/endothelial cell marker CD146. Arrowheads indicate vessels. Scale bar, 100 μm. **g**, Representative fluorescent micrographs of *sgPlxnb2* transduced livers showing nuclear DAPI stain (blue), and “-SMA (green) and periostin (POSTN, red) immunoreactivity. Scale bar, 20 μm. **h**, Quantification of metastatic foci per liver section 1 week after injection of AKPS organoids treated ex vivo with rmPlexin B2 or control. **a, c, h**, Barplots indicate mean ± SD. Dots represent individual mice. Two-tailed unpaired t-test.

**Extended Data Figure 5.**
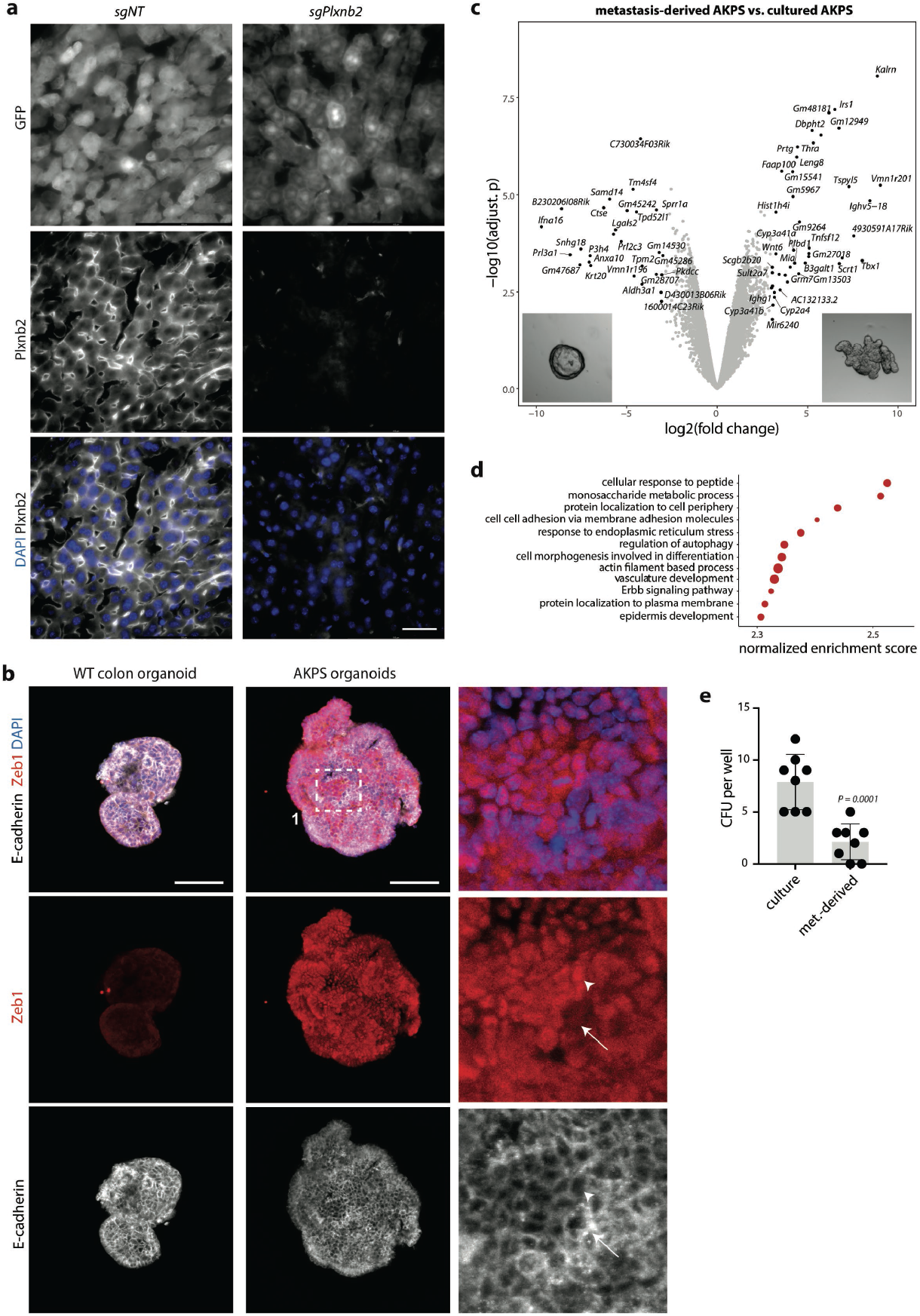
*Plxnb2*-overexpression in hepatocytes promotes metastatic seeding in vivo. **a**, Representative fluorescent micrographs of *AAV8-U6-sgNT-EF1A-EGFP* (*sgNT*) or *AAV8-U6-sgPlxnb2-EF1A-EGFP* (*sgPlnxb2*) transfected livers of *AlbCre;Cas9* mice showing nuclear DAPI staining, endogenous GFP signal and Plexin B2 immunoreactivity. Scale bar, 100 μm. **b**, Representative fluorescent micrographs E-cadherin and Zeb1 staining in WT colon organoids and AKPS organoids. Arrow shows E-cadherin^high^Zeb1^low^ cell, arrowhead indicates E-cadherin^low^Zeb1^high^. **c**, Differentially expressed genes between AKPS-organoid lines (n = 2) derived from metastases and AKPS-organoid lines (n = 3) kept only in culture. Labels indicate significantly top differentially regulated genes. Representative micrographs shown at the bottom. **d**, GO-terms enriched in differentially expressed genes in AKPS lines derived from metastases. **e**, CFU per well upon single-cell seeding of AKPS organoids kept in culture (culture) or derived from metastases (met.-derived). Barplots indicate mean ± SD. Dots represent individual wells, two-tailed unpaired t-test.

**Extended Data Figure 6.**
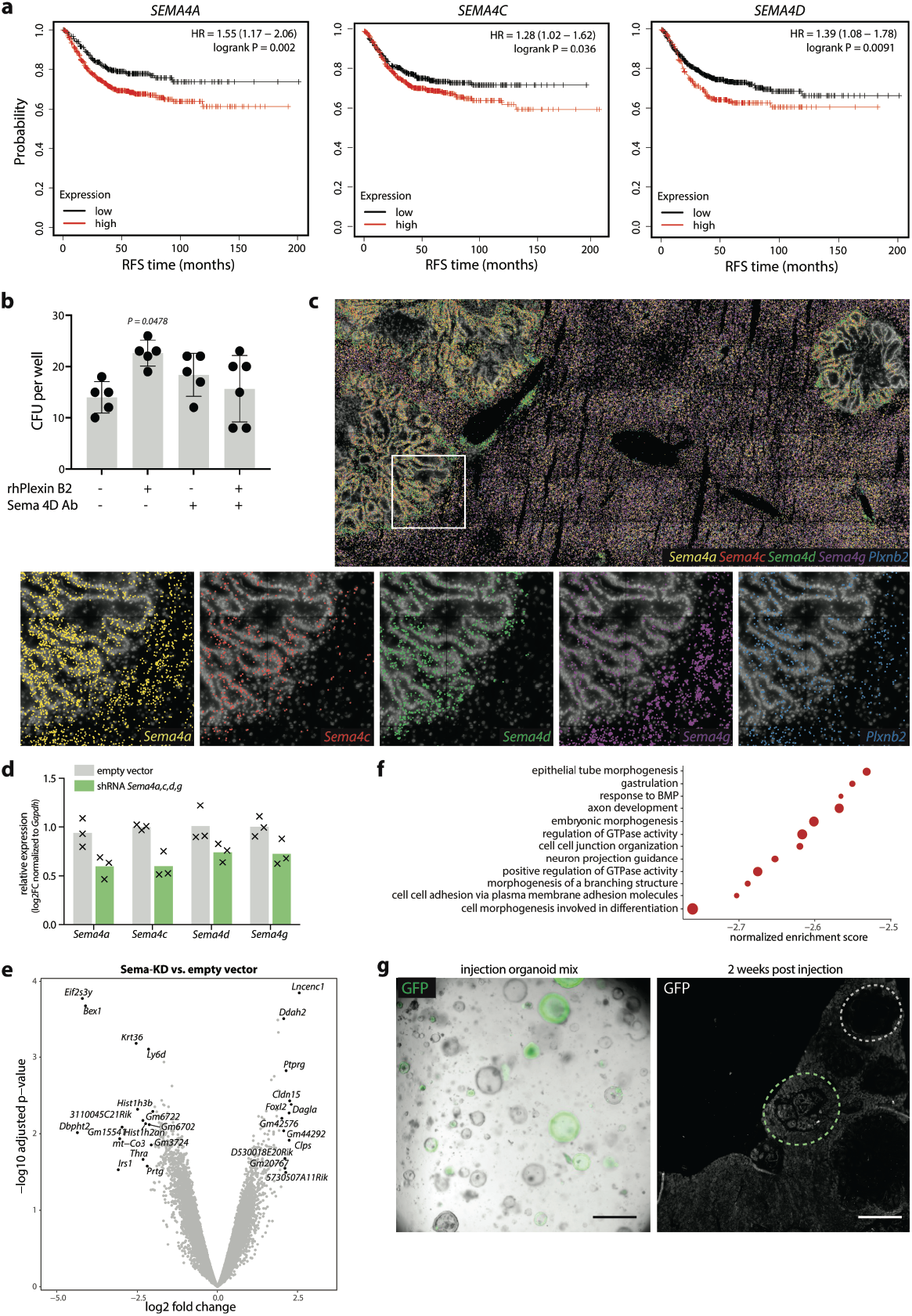
Plexin B2 promotes mesenchymal-to-epithelial transition. **a**, Kaplan-Meier analysis (http://kmplot.com) of the recurrence free survival (RFS) of CRC patients stratified by *SEMA4A, SEMA4C* and *SEMA4D* expression. n = 1211 patients. **b**, CFU per well of patient-derived CRC organoids (PDOs) cultured with 2 μg/mL rhPlexin B2 and 1 ng/μL anti-Sema4D antibody, or IgG4 isotype control. Ordinary one-way ANOVA, comparing each treatment group to untreated control. Tukey’s correction for multiple testing. **c**, Multiplexed in situ hybridization of *Sema4a, 4c, 4d, 4g* and *Plxnb2* transcripts in AKPS metastases. **d**, Gene expression of class 4 semaphorins in AKPS organoids harboring shRNAs targeting *Sema4a, Sema4c, Sema4d* and Sema4g, relative to their expression organoids harboring empty vectors. Log2FC normalized to Gapdh. Crosses indicate technical replicates. **e**, *Differentially* expressed genes between Sema-KD organoids (n = 3) and control empty vector organoids (n = 3). Labels indicate significantly top differentially regulated genes. Representative micrographs shown at the bottom. **f**, GO-terms depleted in differentially expressed genes in Sema-KD organoids. **g**, Left, representative flourescent micrographs of pre-injection 1:1 mixed cultures of Sema-KD AKPS organoids (GFP^+^) and control organoids (empty vector, GFP^-^). Scale bar, 100 μm. Right, representative GFP immunoreactivity in livers co-injected with Sema-KD and control organoids, which form GFP^+^ metastases (example indicated by green dashed line) and GFP^-^ metastases (example indicated by gray dashed line). Scale bar, 200 μm.

## Supplemental Gating Strategy

**Supplementary Table S1: sgRNA library**.

**Supplementary Table S2: sequences of primers, sgRNAs and smFISH probes**.

