## Supplementary figures and images for "In vivo screening of tumor-hepatocyte interactions identifies Plexin B2 as a gatekeeper of liver metastasis"

### Supplemental Gating Strategy

#### Supplemental Gating Strategy

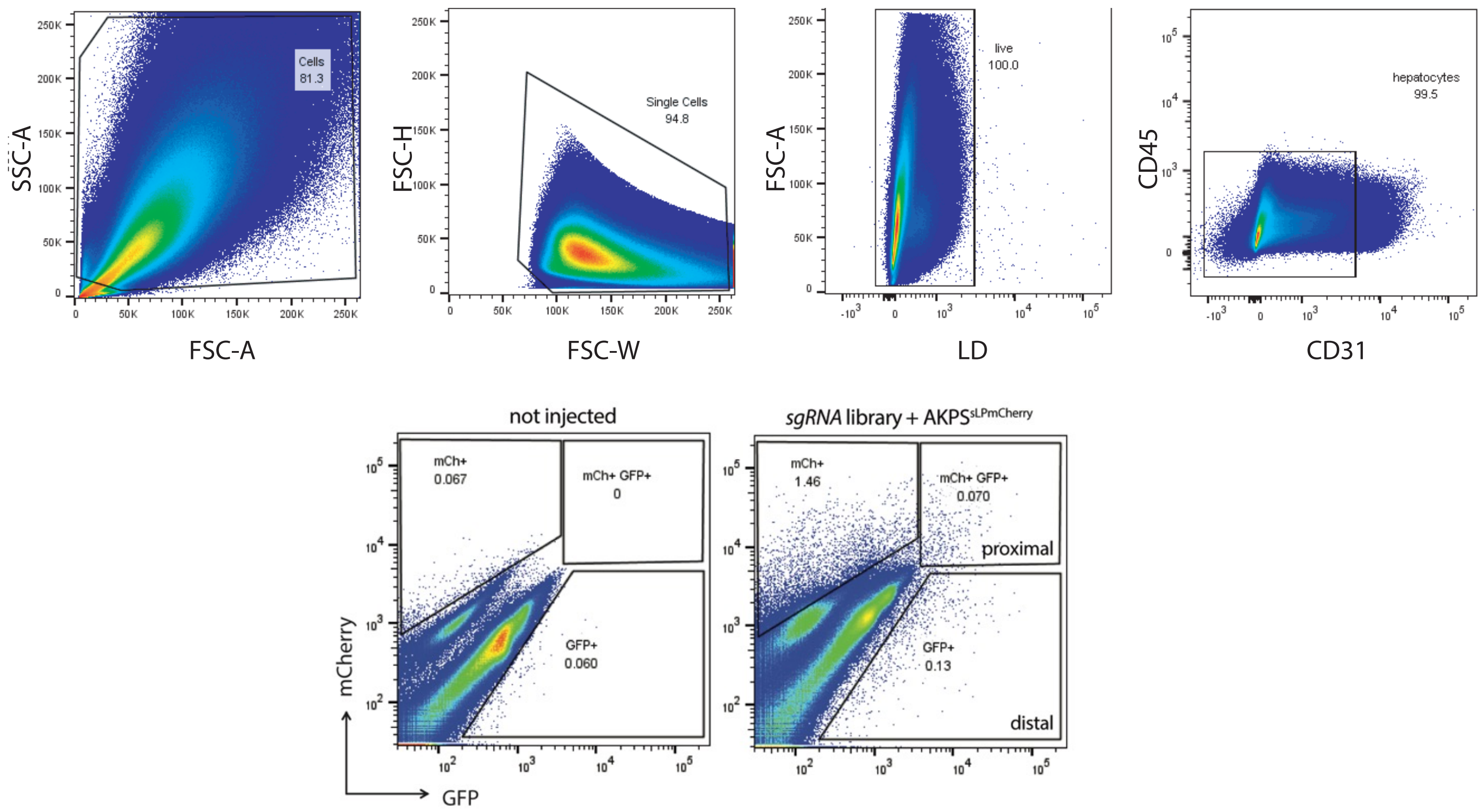
